# Depolarization of astrocytes in the basolateral amygdala restores WFS1 neuronal activity and rescues impaired risk-avoidance behavior in *DISC1^TM^* mice

**DOI:** 10.1101/2022.08.28.505618

**Authors:** Xinyi Zhou, Qian Xiao, Shuai Chen, Xirong Xu, Yuchuan Hong, Yuewen Chen, Liping Wang, Yu Chen, Fan Yang, Jie Tu

## Abstract

Many mental illnesses are accompanied by abnormal risk-avoidance behavior, yet we have only a limited understanding of the neuronal regulatory mechanisms involved. We previously established an inducible *DISC1-N* terminal fragment transgenic mouse (*DISC1-N^TM^*) model which has exhibited risk-avoidance deficiency. Using this model, we analyzed differentially expressed genes (DEGs) using snRNA-seq and the results indicate impaired neuron-astrocyte interactions. We used optogenetic tools to modulate astrocytes in the basolateral amygdala (BLA) and found that ChR2-expressing astrocytes were able to rescue risk-avoidance impairment in *DISC1-N^TM^* mice. Using patch clamp recordings combined with signal-cell qPCR, we found impaired excitability of BLA^WFS1^ neurons in *DISC1-N^TM^* mice and that ChR2-expressing astrocytes can induce action potentials (APs) in WFS1 neurons, which restores WFS1 neuronal activity. WFS1 neurons are necessary for BLA astrocytes to modulate impaired risk-avoidance behavior. These finding provide new insights into mechanisms of astrocyte-neuron interactions and suggest that BLA astrocytes may be a promising target for impaired risk avoidance in mental illness.

**Highlights:** ChR2-expressing astrocytes in the BLA rescue impaired risk-avoidance behavior in *DISC1-N^TM^* mice.

Astrocytes in the BLA modulate different nearby neurons to different degrees. Depolarization of BLA astrocytes restore neuronal activity in WFS1 neurons. Astrocytes in the BLA modulate WFS1 neurons via NMDARs.

## Introduction

Mental illnesses carry some of the greatest burdens among all diseases (Vigo et al., 2016). According to statistics, in most countries in the world, one-third of the population has suffered from at least one mental illness at some time in their life (2000). Mental illnesses have complex symptomatic phenotypes and around half of are accompanied by abnormal risk-avoidance behavior (Lorian and Grisham, 2011, 2012; Maner and Schmidt, 2006; Reddy et al., 2014). For example, patients with anxiety disorders, depression and autism exhibit high risk-avoidance behaviors (Allen and Badcock, 2003; South et al., 2014). Conversely, patient with mental illnesses such as substance-use disorder and borderline personality disorder show reduced risk avoidance (Endrass et al., 2016; Grant et al., 2000; Yamamoto et al., 2015). Mental illnesses have complex pathogenesis and there is a lack of effective assessment methods. Exploring the neural mechanisms underlying abnormal risk-avoidance behavior can provide new insights to our understanding of mental-illness pathogenesis and help to find promising trans-diagnostic factors and therapeutic targets.

The *Disrupted-in-schizophrenia 1* (*DISC1*) gene was initially reported in a translocation mutation segregating with major mental illnesses in a Scottish family (Millar et al., 2000). The *DISC1* gene is thought to be a genetic risk factor for a spectrum of major mental illness (Porteous et al., 2011), including schizophrenia (Porteous et al., 2014), bipolar disorder (BP) (Hennah et al., 2009) and major depressive disorder (MDD) (Duff et al., 2013; Hashimoto et al., 2006). Different types of *DISC1* transgenic mice have already been constructed to study phenotypes found in mental illnesses, including anxiety-like behavior (Wang et al., 2019), depressive-like behavior (Gómez-Sintes et al., 2014; Hikida et al., 2007), abnormal social behavior (Kaminitz et al., 2014), and impaired learning and memory (Wang et al., 2019). However, risk-avoidance behavior, a phenotype shared by multiple mental illnesses, has not been well studied in *DISC1* animal models. Our group previously used an inducible *DISC1-N* terminal fragment transgenic mouse model and reported that it has reduced risk-avoidance behavior (Zhou et al., 2021). Here, we further explore the neural mechanisms underlying this risk-avoidance behavior in *DISC1-N^TM^ mice*.

Neuroimaging data show that the amygdala plays a key role in calculating the expected value of loss-related outcomes (Yacubian et al., 2006). Studies in rats have found that neural circuits involving the BLA are critical for active avoidance behavior (Diehl et al., 2020; Ramirez et al., 2015). The BLA, as the main afferent input nucleus in the amygdala, is a core nucleus that regulates defensive behaviors and fear emotions (Stujenske and Likhtik, 2017; Tovote et al., 2015). Fear of specific stimuli leads to risk-avoidance behavior (Eaton et al., 2018), which is a type of defensive behavior (Hubbard et al., 2004). Therefore, structural or functional abnormalities of cells in the BLA may lead to abnormal risk-avoidance behavior. In studies that target the BLA, astrocytes are typically investigated in addition to neurons. Astrocytes play a critical role in the maintenance of neuronal health and function (Jha et al., 2018). Impairments of astrocyte function have been implicated in various mental illnesses, including depression disorders (Stujenske and Likhtik, 2017), schizophrenia (Hubbard et al., 2004) and bipolar disorder (Toker et al., 2018). However, the function of astrocyte-neuron communications in risk avoidance behavior related to mental illnesses has not been fully investigated.

Our study shows that communication between astrocytes and neurons within the BLA of *DISC1-N^TM^* mice is impaired. Depolarization of BLA astrocytes directly acts on WFS1 neurons via N-methyl-D-aspartate receptors (NMDARs) to rescue the abnormal risk-avoidance behavior found in *DISC1-N^TM^* mice.

## Results

### *DISC1-N^TM^* mice exhibit impaired risk-avoidance behavior

Paradigms used to test risk-avoidance behavior are based on approach-avoidance conflict, in which animals experience opposing impulses of desire and fear (Blomeley et al., 2018; Millan, 2003). Exploratory conflict paradigms are based on conflict between the innate drive to explore novel environments and the fear of potential unknown threats (Kirlic et al., 2017a). To obtain a greater insight into the risk-avoidance behavior of *DISC1-N^TM^* mice, we used three exploratory conflict paradigms: the open-field test (OFT), the EPM and a light-dark box (LDB). Tamoxifen (i.p.) was administered to all mice, including wild-type (WT) mice as a control groups, 6 h before testing. We found less risk avoidance in *DISC1-N^TM^* mice compared to the control groups. This was indexed in the OFT by more entries to the center and by more time spent in the center (Figure 1A-C) and in the EPM by more entries to the open arms and more time spent in the open arms (Figure 1E-G). It is thought that the center of an open arena and the open arms of the EPM are perceived by mice to be less safe, or risky, areas. There was no significant difference in the mean speed between these two groups during the tests (Figure 1D, H). In the LDB experiment (Figure 1I), *DISC1-N^TM^* mice spent significantly more time in the light compartment compared to control mice (Figure 1J), while there was no difference in the total number of transitions between the two compartments (Figure 1K). These results indicate that *DISC1-N^TM^* mice have impaired risk-avoidance behavior.

**Figure 1.**
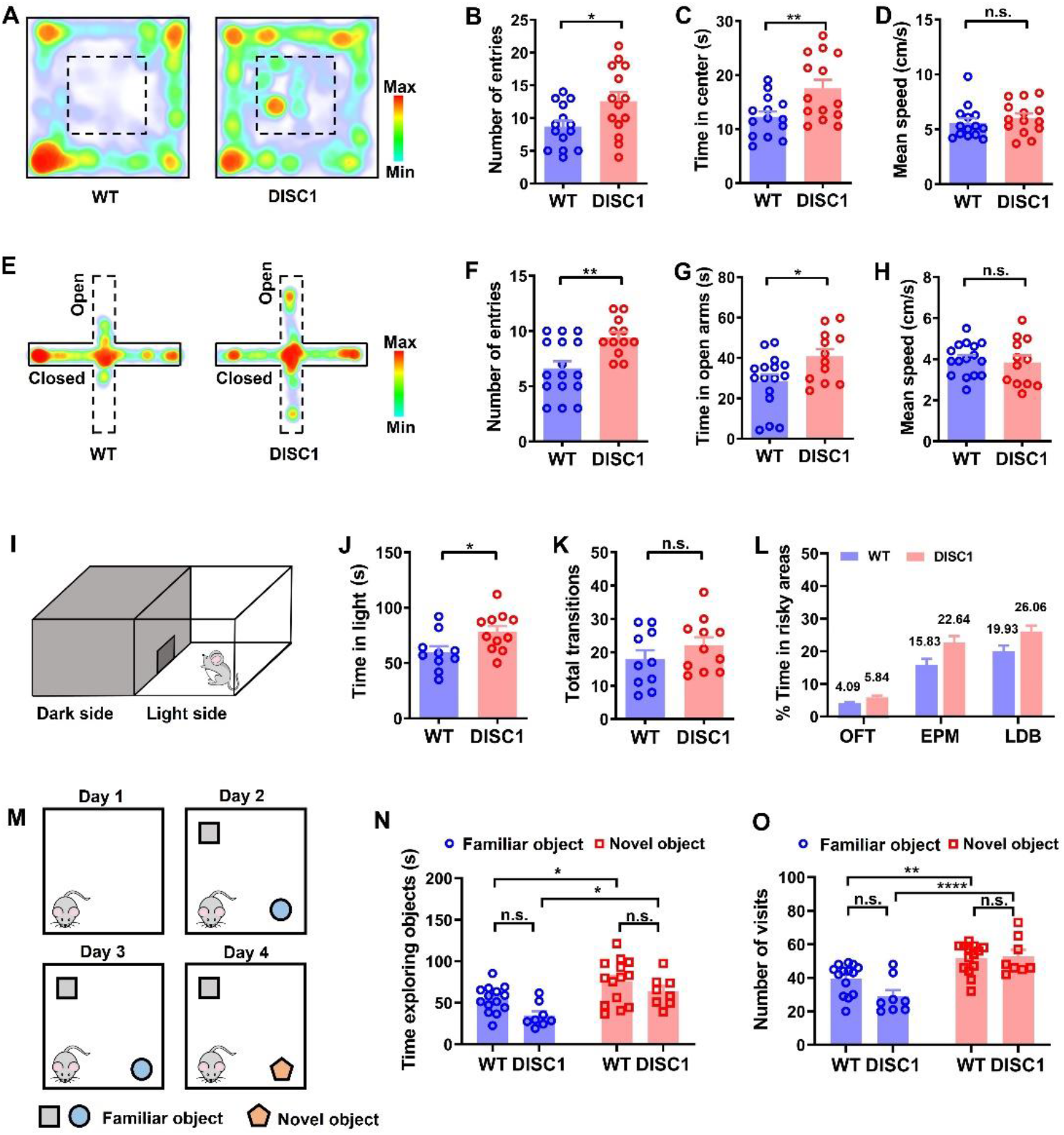
*DISC1-N^TM^* mice have impaired risk-avoidance behavior. (A) Example heatmap showing an OFT in *DISC1-N^TM^* and WT mice. (B-D) The number of entries to the center area (B), time spent in the center area (C), and mean speed (D) during the OFT (unpaired t-test, \**P* = 0.0283, \*\**P* = 0.0096; n _(WT)_ =14, n _(DISC1)_ =14). (E) Example heatmap showing the EPM test in *DISC1-N^TM^* and WT mice. (F-H) The number of entries to the open arms (F), time spent in the open arms (G) and mean speed (H) during the EPM test (unpaired t-test, \**P* = 0.0212, \*\**P* = 0.0031; n _(WT)_ =16, n _(DISC1)_ =12). (I) Schematic of the light-dark box. (J-K) Time spent in the light box (J) and the total number of transitions between the light and dark boxes (K) during the light-dark box test (unpaired t-test, \**P* =0.0248; n _(WT)_ =10, n _(DISC1)_ =11). (L) The percentage display of time spent in the risky areas during OFT, EPM and LDB of *DISC1-N^TM^* and WT mice. (M) Schematic of the NOR test. (N-O) The exploration time (N) and the number of visits (O) to objects in the NOR task (two-way ANOVA, \**P* (up) = 0.041, \**P* (down) = 0.0327, \*\**P* = 0.0038, \*\*\*\**P* < 0.001; n _(WT)_ =14, n _(DISC1)_ =8).

To determine whether the results we observed in the exploratory conflict paradigms were influenced by increased novelty seeking, we tested *DISC1-N^TM^* mice on a novel object recognition (NOR) task to determine whether they exhibited a high novelty-seeking phenotype (Figure 1M). The results showed that both *DISC1-N^TM^* and WT control mice explored the novel object more than the familiar objects, indexed by a significantly longer time and had a significantly greater number of visits to the novel object compared to the familiar objects (Figure 1N, O). These results suggest that *DISC1-N^TM^* mice have similar novelty-seeking behavior to WT mice. These results suggest that the reduced risk avoidance in *DISC1-N^TM^* mice is not due to elevated novelty-seeking behavior.

The experimental paradigms used (OFT, EPM and LDB) are based on unknown risks, that is, no threat obvious perception. But how do *DISC1-N^TM^* mice respond when there is a known risk or dangers? To answer this question, we conducted a looming paradigm test of fear responses to visual stimulation mimicking an overhead airborne predator (Zhou et al., 2019) (Figure S1A). Following looming stimuli, *DISC1-N^TM^* mice had no difference in response latency or in time spent returning to the shelter between the two groups (Figure S1C, D). Interestingly, after *DISC1-N^TM^* mice returned to the shelter, they spent significantly less time in the shelter than that of the WT control group (Figure S1B). These results suggest that *DISC1-N^TM^* mice have abnormal predator fear responses and the lower amount of time spent in the shelter further suggests a reduced risk-avoidance phenotype in *DISC1-N^TM^* mice.

In addition to risk-avoidance related behavior, we also tested depressive-like behavior in *DISC1-N^TM^* mice using the sucrose preference test (Figure S1E). We found no significant difference in the total consumption of liquid (Figure S1F) or in the percentage of sucrose water intake (Figure S1G) between *DISC1-N^TM^* mice and control mice following 24-hr water deprivation. This result indicates that *DISC1-N^TM^* mice do not have anhedonia symptoms and we speculate that *DISC1-N^TM^* mice do not have depressive-like behaviors.

### Astrocytes and neurons in the BLA of *DISC1-N^TM^* mice have abnormal interactions

Having found impaired risk-avoidance in *DISC1-N^TM^* mice, we wondered whether BLA neurons have functional abnormalities associated with the reduced risk-avoidance behavior. We investigated the electrophysiological profiles of BLA neurons in *DISC1-N^TM^* and WT mice using whole-cell patch clamp recordings (Figure S2A). Six hours after tamoxifen injection, we prepared acute brain slices and recorded neurons in the BLA under the current clamp model. There was no difference in the resting membrane potential (RMP) of BLA neurons between the two groups (Figure S2B). We then gave a series of step-current stimulation to the neurons from -100 pA to +140 pA (interval: 10pA). Following the increase of current stimulation, neurons began to generate action potentials (APs) (average threshold current: WT= 56pA, *DISC1-N^TM^*= 57pA). We compared the current-induced firing rates and found that neurons in *DISC1-N^TM^* mice had significantly lower firing rates than WT mice following both 130-pA and 140-pA current stimulation (Figure S2C, F, G). This indicates that BLA neurons were less excitable in *DISC1-N^TM^* mice than those in the WT mice. We also compared the amplitude and half-wave width of APs evoked by 140-pA current and found no difference between the two groups (Figure S2D, E).

In a previous investigation of the BLA, we found that different rates of glutamatergic neuron photostimulation (ChR2-expression) led to varying degrees of risk-avoidance behaviors in *DISC1-N^TM^* mice (Zhou et al., 2022). Therefore, we speculated the glutamatergic neurons in the BLA were heterogeneous. Next, we used single-nucleus RNA sequencing (snRNA-seq) to determine BLA cell transcriptomes in *DISC1-N^TM^* mice. We sampled 7171 nuclei from *DISC1-N^TM^* brain samples and 6878 nuclei from WT brain samples. We obtained 12 cell clusters (Figure 2A) and classified them into seven cell types based on their respective transcriptional profiles and previously reported cell-type markers in both *DISC1-N^TM^* and WT mice: astrocytes (*AQP4^+^*, 15.35% vs. 15.46%), endothelial cells (*CLDN5^+^*, 1.26% vs. 0.80%), excitatory neurons (*CAMK2A^+^,* 48.31% vs. 49.48%), inhibitory neurons (*GAD1^+^*, 10.64% vs. 10.69%), microglia (*C1QA^+^*, 6.28% vs. 5.05%), oligodendrocytes (*MBP^+^*, 9.69% vs. 9.19%) and oligodendrocyte precursor cells (*PDGFRA^+^* 8.48% vs. 9.35%) (Figure 2B). We counted the expression of differentially expressed genes (DEGs) in *DISC1-N^TM^* and WT samples in different cell types. Excitatory neurons had 166 DEGs, which was the highest number found, followed by astrocytes with 40 DEGs (Figure S3A). Among 249 DEGs found, only 1 was differentially expressed in all cell types (Figure S3B), suggesting that most observed *DISC1-N^TM^*-associated transcriptomic changes are cell-type specific.

**Figure 2.**
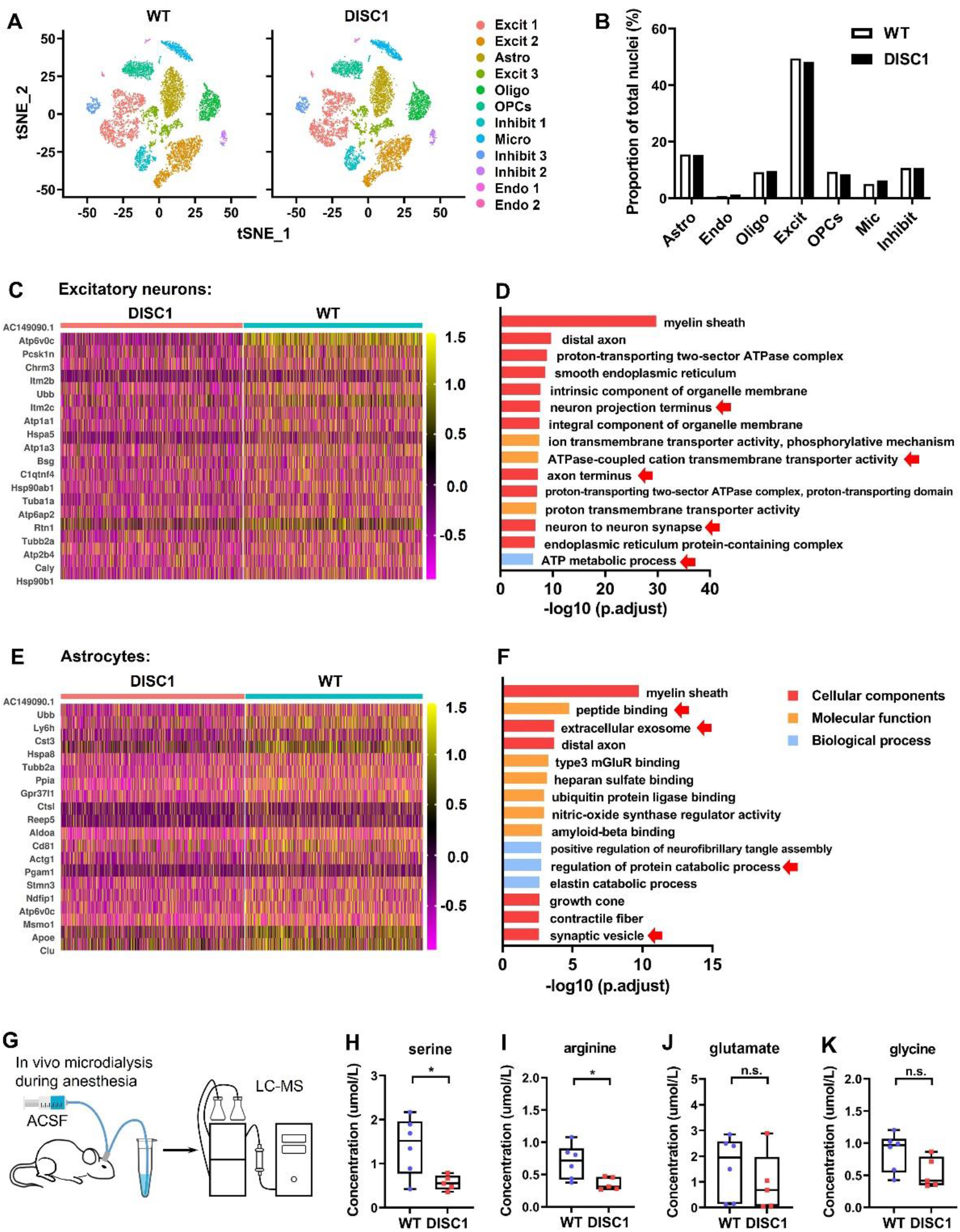
snRNA-seq analysis of BLA cells in WT and *DISC1-N^TM^* mice. (A-B) t-SNE plots (A) and bar plot (B) showing the proportions of the seven major cell types found in the WT and *DISC1-N^TM^*BLA samples. (C-D) Heatmap showing the expression levels of the most enriched genes in the *DISC1-N^TM^*-down-regulated astrocytes (C) and excitatory neurons (D) (adjusted P < 0.1, log2 fold change ≤ −0.1). (E-F) GO analysis of the down-regulated transcriptomic signature of astrocytes (E) and excitatory neurons (F) (the top 15 most relevant indicators are shown). (G) Schematic showing the microdialysis experiment. (H-K) The partial results of LC-MS analysis of the dialysate from the BLA of WT and *DISC1-N^TM^* mice (unpaired t-test, from left to right: \**P* = 0.0223 and 0.0258; n _(WT)_ =6, n _(DISC1)_ =5).

Next, to determine the DEGs and gene product enrichment, we performed Gene Ontology for down-regulated DEGs in neurons and astrocytes (neurons and astrocytes were enriched with down-regulated signature genes). We found that, in the BLA of *DISC1-N^TM^* mice, excitatory neurons had abnormal gene expression profiles which were highly enriched in genes related to ATP metabolism, myelin sheath, axon terminals, synapses, transmembrane transporters and other functions (Figure 2C). These enrichments suggest that the interactions between excitatory neurons and other cells may be affected. The transcriptome profile of the DISC1-down-regulated astrocytes subpopulation was characterized by enriched expression of genes associated with protein catabolism processes, receptor aggregation, exosomes, vesicles, and glutamate receptor binding (Figure 2E). These results suggest that astrocytes in *DISC1-N^TM^* mice had structural or functional abnormalities related to receiving external signal stimulation and in transmitting signals through the release of glial transmitters from vesicles.

To determine whether BLA astrocytes had dysfunctional release of gliotransmitters, we used microdialysis to obtain intercellular material from the BLA from both *DISC1-N^TM^* mice and WT mice 6 hr after tamoxifen injection. Using liquid chromatography-mass spectrometry (LC-MS), we found a reduction in several amino acids released from cells in the BLA of *DISC1-N^TM^* mice, including serine, arginine, taurine, leucine, and alanine/sarcosine (Figure 2H, I and Figure S3). There was no significant difference in the concentrations of amino acids, such as glutamate, glycine, proline, isoleucine, glutamine, lysine and valine (Figure 2J, K and Figure S3). The decrease in serine concentration indicates that *DISC1-N^TM^* mice have an abnormality in the secretion of D-serine, which plays an important role in regulating physiological activity in neurons. Astrocytes are one of the main sources of D-serine in the central nervous system (CNS), and can release D-serine to regulate neuronal function (Panatier et al., 2006; Papouin et al., 2017a). This, combined with our snRNA-seq results, implies that glial vesicle release may be impaired (lower expression of related DEGs included Calm2, Calm1, Atp6v0c and Hspa8) in BLA astrocytes from *DISC1-N^TM^* mice. We speculate that the interaction between astrocytes and neurons in *DISC1-N^TM^* mice is abnormal, which in turn affects the function of excitatory neurons in this brain region, resulting in the behavioral manifestations of reduced risk avoidance in mice.

### Risk-avoidance impairment in *DISC1-N^TM^* mice can be rescued by photostimulation of BLA astrocytes

To test the hypothesis that astrocytes in the BLA of *DISC1-N^TM^* mice have abnormal astrocyte-neuron interactions, we used optogenetic tools to modulate astrocytes and determine whether impaired risk avoidance in *DISC1-N^TM^* mice could be rescued. An AAV vector encoding ChR2-mCherry under the control of the GFAP promotor was used to transfect BLA astrocytes (Figure 3A). To confirm the specificity of the AAV-GFAP-ChR2-mCherry virus, we performed immunohistochemistry staining of S100 (astrocyte-specific expressing markers) on brain slices expressing virus (Figure 3C). The co-expression of S100 with mCherry reached 88.69%, while the co-expression of NeuN with mCherry was only 2.42% (Figure 3B-C). The immunohistochemistry results indicated that the GFAP promoter carried by adenovirus specific expressed ChR2 on BLA astrocytes.

**Figure 3.**
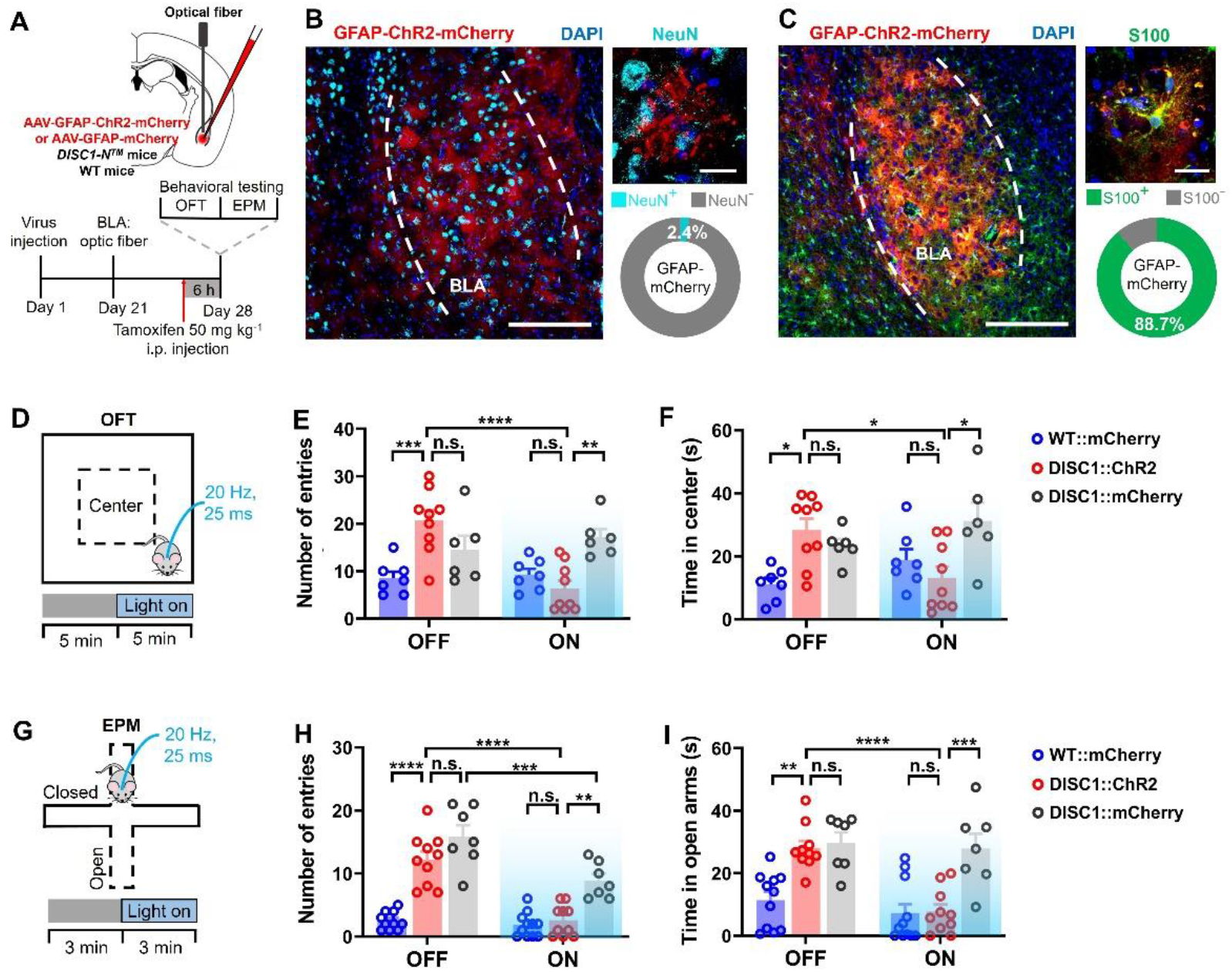
Optogenetic stimulation of BLA astrocytes rescues risk-avoidance impairment in *DISC1-N^TM^* mice. (A) Schematic showing modulation of astrocytes in the BLA using optogenetic tools. (B-C) Representative images and percentage statistics showing the co-expression of ChR2-expressing cells with either NeuN-expressing cells (cyan, B) or S100-expressing cells (green, C) in the BLA (n _(DISC1)_ =3). (D) Schematic showing the OFT paradigm. (E-F) The number of entries to (E) and time spent in the center (F) before and after blue-light (470 nm) stimulation during the OFT (two-way ANOVA, left: \*\**P* =0.0045, \*\*\**P* =0.0007, \*\*\*\**P* < 0.0001, right: \**P* =0.0157, 0.0261 and 0.0144 from left to right; n _(WT-mCherry)_ =7, n _(DISC1-ChR2)_ =9, n _(DISC1-mCherry)_ =6). (G) Schematic showing the EPM test paradigm. (H-I) The number of entries to (H) and time spent in the open arms (I) before and after blue-light (470 nm) stimulation during the EPM test (two-way ANOVA, left: \*\**P* =0.0013, \*\*\**P* = 0.0009, \*\*\*\**P* < 0.0001, right: \*\**P* = 0.0013, \*\*\**P* = 0.0005, \*\*\*\**P* < 0.0001; n _(WT-mCherry)_ =11, n _(DISC1-ChR2)_ =10, n _(DISC1-mCherry)_ =7).

The OFT and EPM paradigms were used to examine the impact of astrocyte manipulation on behavioral performance in *DISC1-N^TM^* mice. Six hours after tamoxifen injections, mice were placed in either the open field or the EPM and were allowed to explore freely for a fixed period of time (“light-off” stage). Then, blue light stimulation (25 ms per pulse, 20 Hz) was directly shone on the BLA region for the same duration (“light-on” stage) (Figure 3D, G).

In addition to *DISC1-N^TM^* mice that unilaterally express AAV9-GFAP-ChR2-mCherry in the BLA, we also set up two control groups: WT mice and *DISC1-N^TM^* mice unilaterally expressing AAV9-GFAP-mCherry in the BLA. During the light-off stage, *DISC1-N^TM^* mice spent a significantly longer time in the risky areas of either maze (center of open field and open arms in the EPM) and had a significantly larger number of entries into each risky area compared to the WT group (Figure 3E, F, H, I). During the light-on phase, ChR2-expressing *DISC1-N^TM^* mice not only spent less time in and had fewer entries to the risky areas compared to light-off phase, but also reached the same level of behavior as WT mice during light-on stage (Figure 3E, F, H, I). This was in contrast to the nonexpressing-ChR2 *DISC1-N^TM^* mice, which maintained their reduced risk-avoidance behavior (compared with the ChR2-expressing *DISC1-N^TM^* mice) during the light-on stage (Figure 3E, F, H, I). The results of nonexpressing-ChR2 *DISC1-N^TM^* mice suggest that there is no rescue of impaired risk-avoidance behavior in *DISC1-N^TM^* mice unless astrocytes express ChR2 during the light-on stage. This excludes the possibility that the interference of the blue laser stimulation affects behavioral output, and the possibility that the altered behavioral outputs on the OFT and EPM are a result of habituation to the experimental paradigms. In conclusion, the behavioral results of both the OFT and EPM test indicate that optogenetic tools used to modulate ChR2-expressing astrocytes in the BLA can rescue risk-avoidance impairment in *DISC1-N^TM^* mice.

We then employed a pharmacogenetic approach to examine whether astrocytes in the BLA can rescue risk-avoidance impairment in *DISC1-N^TM^* mice. We use the same mouse groups as with the astrocyte-modulating optogenetic experiment above, except using *DISC1-N^TM^* mice expressing AAV9-GFAP-hM3Dq-mCherry in the BLA rather than AAV9-GFAP-ChR2-mCherry (Figure S5D). Mice in all groups were intraperitoneally injected with CNO 30 min before the start of the experiment. In both the OFT and the EPM, the time spent in the risky areas and the number of entries to the risky areas of hM3Dq-expressing *DISC1-N^TM^* mice were at the same level as the WT control group and significantly lower than the nonexpressing-hM3Dq *DISC1-N^TM^* mice (Figure S5G, H, J, K). These experimental results are consistent with the behavioral results using optogenetic techniques to modulate astrocytes (Figure 3D-I), and there was no difference in the average mean speed between any of the three groups in the behavioral experiments (Figure S5I, L). Both optogenetic and pharmacogenetic experimental results suggest that the risk-avoidance impairment in *DISC1-N^TM^* mice can be rescued by modulating astrocytes in the BLA.

### BLA Neurons exhibited distinct electrophysiological responses to photostimulated astrocytes

The behavioral tests confirmed that modulation of astrocytes can rescue risk-avoidance impairment in *DISC1-N^TM^* mice, but we still did not know how astrocytes affect neurons. To approach this, we used patch clamp recording to explore the electrophysiological mechanisms underlying the regulation of impaired risk-avoidance behavior in astrocytes. Coronal brain slices containing the BLA were obtained from WT mice 6 hours after tamoxifen injections. ChR2-expressing astrocytes were photostimulated when recording the electrophysiological activity of nearby neurons (Figure 4A) (the distance between ChR2-expressing astrocytes and recorded neurons was less than 100 μm, Figure S6A). When 15 s of blue light stimulation (25 ms per pulse, 20 Hz) was shone onto the surface of the brain slice, we recorded two different electrophysiological responses from the nearby neurons. The membrane potential of one neuronal response was a depolarization, and eventually generation of APs, which we named high-response neurons (Figure 4B). In another set of neurons, the membrane potential was slightly raised followed by either a return to baseline or no further rise (average membrane potential rise: 5.16mV), and these neurons, which we named low-response neurons, failed to produce APs during the blue-light stimulation period (Figure 4C). We speculated that these neurons are two different neuronal subtypes in the BLA with different electrophysiological responses to ChR2-expressing astrocytes. We measured the time from onset of blue-light stimulation to when high-response neurons produced APs and it took an average of more than 3 seconds (3.60 ± 2.40 s, Figure 4D), which was much longer than the time required for synaptic signaling between neurons.

**Figure 4.**
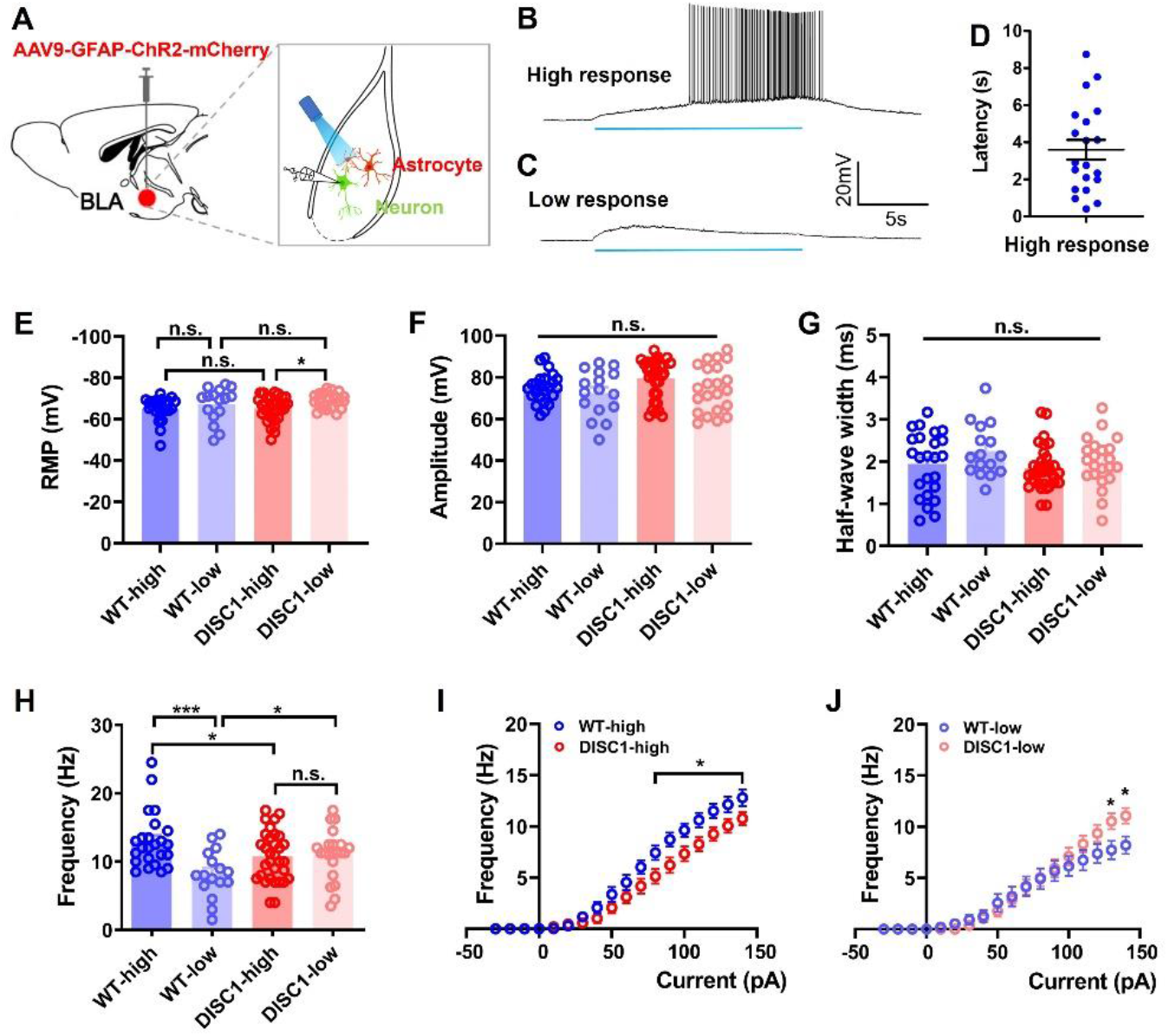
Astrocytes in the BLA differentially modulate different nearby neurons. (A) Schematic showing patch clamp recording of neurons in the BLA. (B-C) High-response (B) and low-response neurons (C) recorded by patch clamp during photostimulation. (D) The latency of high-response neurons from onset of light to generation of APs (n = 20 neurons from 14 mice). (E) Comparison of the RMP of high-response and low-response neurons in *DISC1-N^TM^* mice and WT mice (unpaired t-test, \**P* = 0.0117; n _(WT-high)_ = 24 neurons from 17 mice, n _(WT-low)_ =16 neurons from 9 mice, n _(DISC1-high)_ = 32 neurons from 23 mice, n _(DISC1-low)_ =22 neurons from 15 mice). (F-H) Comparison of the amplitude (F), half-wave width (G) and frequency (H) of high-response and low-response neurons in *DISC1-N^TM^* mice and WT mice during 140 pA current stimulation (unpaired t-test, \*\*\**P* = 0.0004, left: \**P* = 0.036, right: \**P* = 0.0187; n _(WT-high)_ = 24 neurons from 17 mice, n _(WT-low)_ =16 neurons from 9 mice, n _(DISC1-high)_ = 32 neurons from 23 mice, n _(DISC1-low)_ =22 neurons from 15 mice). (I-J) The I-frequency curve of high-response (I) and low-response neurons (J) in *DISC1-N^TM^* and WT mice (unpaired t-test, \**P* < 0.05; n (WT-high) = 24 neurons from 17 mice, n (WT-low) =16 neurons from 9 mice, n (DISC1-high) = 32 neurons from 23 mice, n (DISC1-low) =22 neurons from 15 mice).

To determine whether blue light exposure caused depolarization of the neuronal membrane potential, we set up a control group in which AAV9-GFAP-mCherry virus was injected into the BLA (Figure S6G). We found that, without the expression of ChR2 in BLA astrocytes, the membrane potentials barely changed when exposed to blue light, which caused neither depolarization nor generation of APs (Figure S6H). These results confirm that the electrophysiological responses of low-response neurons were caused by opto-stimulation of ChR2-expressing astrocytes rather than exposure to blue light. Next, we tested whether different light stimulation patterns resulted in different responses in low-response neurons. We set up different light stimulation patterns with different frequencies or durations. When the blue light stimulation duration was unchanged for a 15-s period during which the frequency was increased from 1 Hz to 40 Hz (pulse width, 25 ms; maximum stimulation frequency, 40 Hz), the membrane potential of low-response neurons became gradually depolarized, reaching a maximum depolarization level of approximately 30 Hz (Figure S6B, E). The depolarization amplitude during the whole process did not exceed 8 mV (Figure S6E). When the stimulation frequency was unchanged and the stimulation duration was increased, the depolarization of the membrane potential gradually stabilized but did not reach the firing threshold (Figure S6C, D, F). These results indicate that the neurons failed to generate APs (when ChR2-expressing astrocytes received photo-stimulation), independent of the duration and frequency of blue-light stimulation.

We also recorded the two different responses of neurons in the *DISC1-N^TM^* mice and calculated the electrophysiological characteristics of these four groups of neurons: high-response neurons in *DISC1-N^TM^* mice (DISC1-high), low-response neurons in *DISC1-N^TM^* mice (DISC1-low), high-response neurons of WT mice (WT-high), and low-response neurons of WT mice (WT-low). We found that DISC1-low neurons had a significantly higher RMP than DISC1-high neurons, whilst there was no significant difference in RMP between WT-low and WT-high neurons (Figure 4E). In current-clamp recording mode, we applied depolarizing step currents (from -100pA to +140pA, 2s duration, 10pA step) to the neurons. We calculated the amplitude, half-wave width and firing frequency of AP produced by neurons under 140 pA current stimulation. There was no difference in the amplitude and half-wave width between the four groups (Figure 4F, G). In a comparison of firing frequency, DISC1-high neurons had lower frequency than WT-high neurons, while DISC1-low neurons had higher frequency than WT-low neurons (Figure 4H). The current-frequency curves show that the significant difference in firing frequency between DISC1-high neurons and WT-high neurons started at 80 pA, whilst the difference between DISC1-low neurons and WT-low neurons started at 130 pA (Figure 4I, J). These results showed that, compared with WT mice, high-response neurons in *DISC1-N^TM^* mice had a lower level of excitability, while low-response neurons had a higher level of excitability. This evidence suggests that the reduced excitability of high-response neurons, which can be regulated by astrocytes and generate APs, is the reason for risk-avoidance impairment in *DISC1-N^TM^* mice.

### Astrocytes in the BLA modulate risk-avoidance behavior through WFS1 neurons

After classifying high- and low-response neurons in the BLA by electrophysiological response to ChR2-expressing astrocytes, we further identified the molecular markers for these two types of neurons. We combined brain slice patch-clamp and single-cell qPCR, pulling out neurons recorded in the patch-clamp by applying negative pressure to the glass electrode tip, followed by single-cell qPCR experiment (Figure 5A). We selected four possible molecular markers of neuronal subtypes: *Calb2* (*Calbindin 2*), *Htr2C* (*5-Hydroxytryptamine Receptor 2C*), *Map4K3* (*Mitogen-Activated Protein Kinase Kinase Kinase Kinase 3*) and *Wfs1* (*Wolfram Syndrome 1*) for qPCR detection. The four target genes were already found to be expressed in the BLA through snRNA-seq (Figure S7A-D) and the transcribed mRNAs or translation proteins have been reported to be associated with mental illness (Diener et al., 1997; Iwamoto and Kato, 2003; Koido et al., 2005; Konig et al., 2020). Results of qPCR analysis showed that the mRNA expression of Wfs1 in high-response neurons was significantly higher than that in low-response neurons, while there was no significant difference in the mRNA expression of the other three molecular markers (Figure 5B). These results suggests that high-response neurons in the BLA were a WFS1-expressing neuronal subtype. To further confirm that WFS1 neurons in the BLA are excitatory neurons, we detected the co-expression of WFS1 with vGlut (excitatory neuron marker) and GAD (inhibitory neuron marker). We found that WFS1 neurons and vGlut in BLA had a very high co-expression ratio of more than 90% (Figure S7F, G), and a low co-expression ratio with GAD (1.55%, Figure S7F, H). These results showed that the WFS1 neurons in BLA are excitatory neurons.

**Figure 5.**
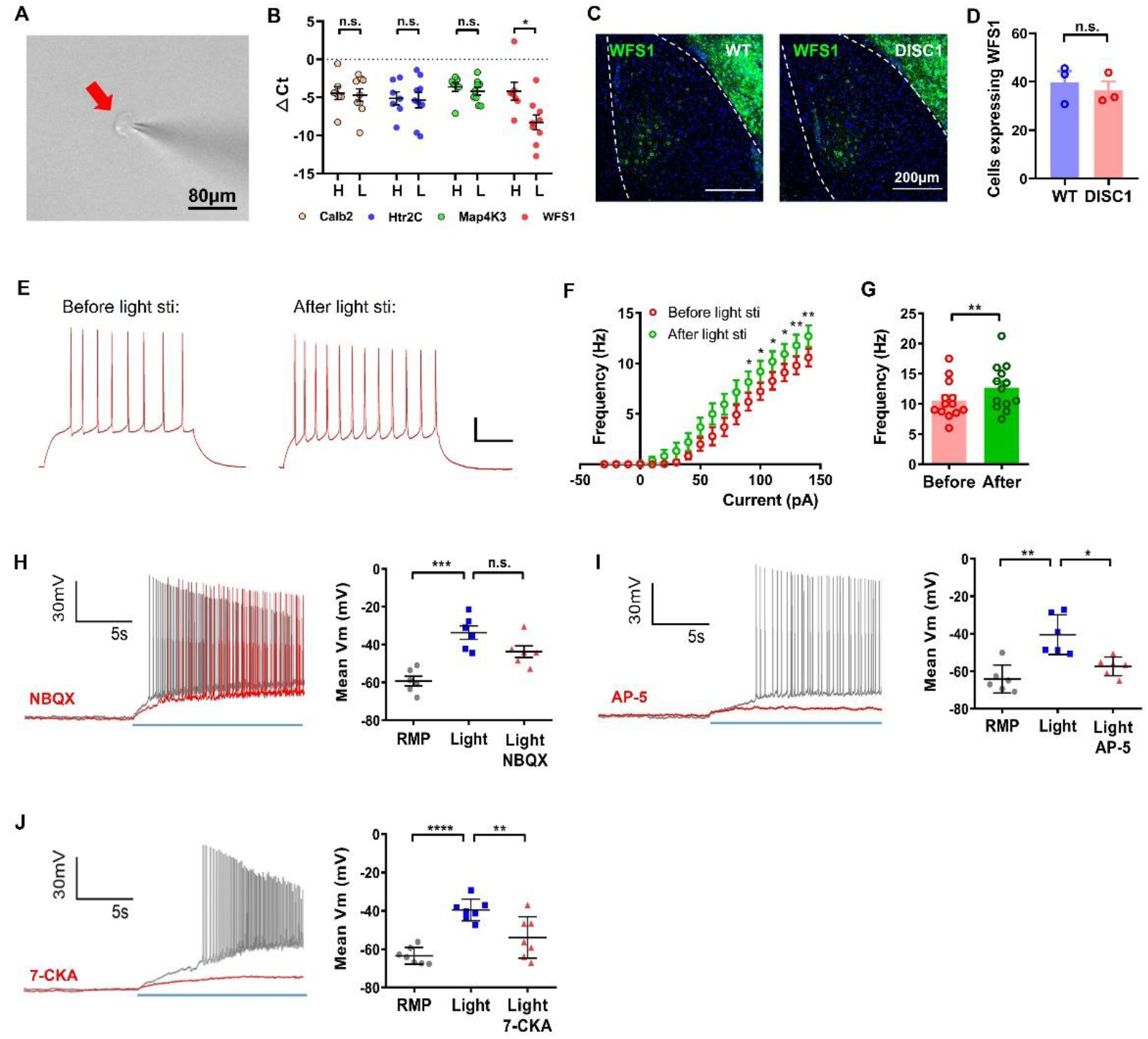
Astrocytes restore WFS1 neuronal activity through NMDARs. (A) Schematic showing a glass pipette pulling a neuron from the surface of a brain slice. (B) The WFS1 marker was highly expressed in high-response neurons (unpaired t-test, \*\**P* = 0.0178; n _(high-response neurons)_ =7 neurons from 7 WT mice, n _(low-response neurons)_ =9 neurons from 6 WT mice). (C) Representative staining of cells in the BLA with WFS1 (green) and DAPI (blue). (D) Quantification of WFS1 neurons in *DISC1-N^TM^* and WT mice (unpaired t-test, n _(WT)_ =3, n _(DISC1)_ =3). (E) Representative traces of high-response neuron evoked by 140 pA current stimulation before and after photostimulation of ChR2-expressing astrocytes in *DISC1-N^TM^* mice. (F-G) The I-frequency curve (F) and frequency comparison (G, induced by 140-pA current stimulation) of high-response neurons in *DISC1-N^TM^* mice before and after photostimulation of ChR2-expressing astrocytes (paired t-test, left: \**P* < 0.05, \*\**P*<0.01, right: \*\**P*= 0.0052; n =13 neurons from 11 *DISC1-N^TM^* mice). (H) Left: representative traces of high-response neuron evoked by photostimulation of astrocytes (grey lines). Application of NBQX failed to block APs (red trace). Right: mean membrane potential of high-response neurons when at RMP, photostimulation of astrocytes and application of NBQX (paired t-test, \*\*\**P* = 0.0002; n = 6 neurons from 3 *DISC1-N^TM^* mice). (I) Left: representative traces of high-response neurons evoked by photostimulation of astrocytes (grey lines). Application of AP-5 blocked APs (red trace). Right: mean membrane potential of high-response neurons when neurons were at RMP, photostimulation of astrocytes and application of AP-5 (paired t-test, \**P* = 0.0247, \*\**P* = 0.0076; n = 6 neurons from 4 *DISC1-N^TM^* mice). (J) Left: representative traces of high-response neuron evoked by photostimulation of astrocytes (grey lines). Application of 7-CKA blocked the APs (red trace). Right: mean membrane potential of high-response neurons when neurons were at RMP, photostimulation of astrocytes and application of 7-CKA (paired t-test, \*\**P* = 0.0023, \*\*\*\**P*<0.0001; n = 7 neurons from 5 *DISC1-N^TM^* mice).

Next, we determined whether there was a difference in the number of WFS1 neurons in the BLA between *DISC1-N^TM^* and WT mice. We performed anti-WFS1 immunostaining experiments on the BLA from *DISC1-N^TM^* and WT mice (Figure 5C). We counted the number of WFS1-expressing neurons and found no difference between *DISC1-N^TM^* mice and WT mice (Figure 5D). We also isolated the brain tissue of BLA region and performed qPCR to compare the mRNA expression of the *Wfs1* gene between *DISC1-N^TM^* and WT mice and found that there was no significant difference (Figure S7E). These results suggest that the reduced risk avoidance in *DISC1-N^TM^* mice is not due to differences in the number of WFS1 neurons in the BLA.

To further explore whether the functional abnormalities of WFS1 neurons could be rescued by astrocytes, we compared the current-induced firing frequency changes of WFS1 neurons before and after photostimulation of astrocytes in *DISC1-N^TM^* mice (Figure 5E). We found that the current-induced firing frequency of WFS1 neurons increased after photostimulation of ChR2-expressing astrocytes (Figure 5F, G). This result further suggests that depolarization of BLA astrocytes can rescue the impaired risk-avoidance behavior in *DISC1-N^TM^* mice by restoring WFS1 neuronal activity.

### Astrocytes influence the activity of high-response neurons via NMDA receptors

We found that opto-stimulation of ChR2-expressing astrocytes caused AP firing in WFS1 neurons, but the mechanism by which astrocytes regulate WFS1 neuronal firing is still unknown. We then determined which glutamate receptors the BLA astrocytes used to regulate the function of the high-response neurons. Neurons in the BLA receive many excitatory glutamatergic inputs and express many glutamate receptors (Tovote et al., 2015). We recorded the high-response neurons using patch clamp and tested the effects of two ionotropic glutamate receptor blockers, AP-5 (NMDA receptor blocker, 50 μM) and NBQX (AMPA receptor blocker, 50 μM). AP-5 blocked AP firing in high-response neurons activated by ChR2-expressing astrocytes (Figure 5I), while NBQX failed to block AP firing (Figure 5H). These results indicate that astrocytes in the BLA regulate nearby high-response neurons through NMDA receptors. Next, we tested the addition of 7-chlorokynurenic acid (7-CKA, 20 μM), a selective NMDA antagonist at the glycine-binding site of the NMDA receptor, and found that it blocked AP firing in the high-response neurons induced by light-stimulated astrocytes (Figure 5J), suggesting that co-activation of the glycine-binding site in NMDA receptors is required for astrocytes in the BLA to regulate the activity of high-response neurons.

### WFS1 neurons in the BLA are required for BLA astrocyte modulation of impaired risk-avoidance behavior

To investigate whether WFS1 neurons are necessary for BLA astrocytes to regulate risk-avoidance behavior in *DISC1-N^TM^* mice, we packaged an AAV virus with *Wfs1* as a promoter to transfect hM4Di onto WFS1 neurons to specifically inhibit the firing of WFS1 neurons during photostimulation of ChR2-expressing astrocytes (Figure 6A). There was 70% co-localization of cells expressing the fluorescent protein carried by the AAV-Wfs1-hM4Di-mCherry virus co-localized with WFS1 cells (Figure 6B, C). The inhibitory effect of CNO on hM4Di-expressing cells was verified by patch clamp; CNO was added to the artificial cerebrospinal fluid (ACSF) led to hyperpolarization of the neuronal membrane potential (Figure 6D, E). We unilaterally injected AAV-GFAP-ChR2-eYFP and AAV-Wfs1-hM4Di-mCherry viruses into the BLA of the *DISC1-N^TM^* mice. Unilateral BLA injection of AAV-GFAP-ChR2-eYFP and AAV-Wfs1-mCherry viruses into the *DISC1-N^TM^* mice served as a control group. Intraperitoneal injection of tamoxifen was given 6 hours before the start of the behavioral experiment, and CNO was injected intraperitoneally 30 minutes before the start of the behavioral experiment. In the OFT (Figure 6F), the number of entries and the time spent in the central area were significantly lower in the control group during light-on stage (Figure 6G, H). In the hM4Di-expressing group, there were no significant difference in either the number of entries or time spent in the central area before and after light stimulation (Figure 6I, J). Similar results were also observed in the EPM test (Figure 6K). Both the number of entries and the time spent in the open arms were significantly lower in the control group during the light-on stage in *DISC1-N^TM^* mice (Figure 6L, M). In the hM4Di-expressing group, there were no significant difference in either the number of entries or time spent in the open arms before and after light stimulation (Figure 6N, O). The behavioral results of both the OFT and the EPM test showed that inhibiting the activity of WFS1 neurons in the BLA blocked the effects of astrocytes on risk-avoidance behavior in *DISC1-N^TM^* mice. These results confirm that WFS1 neurons are necessary for astrocytes in BLA to modulate impaired risk-avoidance behavior in *DISC1-N^TM^* mice.

**Figure 6.**
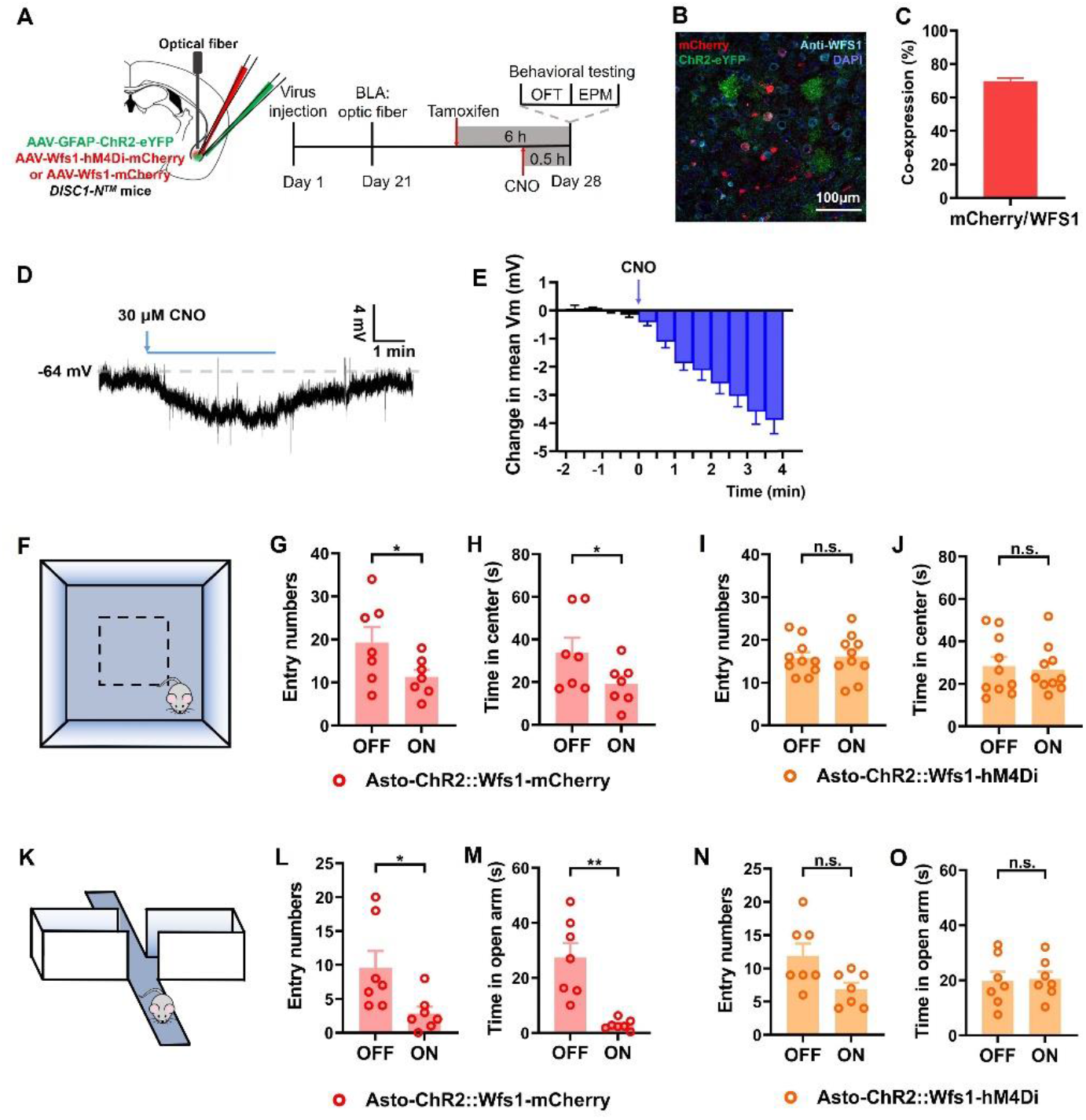
WFS1 neurons are required for astrocyte modulation of risk-avoidance impairment in *DISC1-N^TM^* mice. (A) Schematic showing the experimental timeline. (B) Representative image showing virus expression and anti-WFS1 (cyan) immunofluorescence staining in the BLA (red: AAV-Wfs1-hM4Di-mCherry, green: AAV-GFAP-ChR2-eYFP, blue: DAPI). (C) Quantification of mCherry-expressing cells that simultaneously express WFS1 immunofluorescence staining in the BLA (n (WT) =3). (D) Following CNO administered to brain slices, neurons expressing AAV-Wfs1-hM4Di-mCherry had a decrease in membrane potential. (E) Histogram showing the mean change in membrane potential after CNO administration (n =11 neurons from 5 WT mice). (F) Schematic of the OFT paradigm. (G-H) The number of entries to (G) and time spent in the center (H) of the non-hM4Di-expressing control group during the OFT (paired t-test, number of entries \**P* =0.0386, time spent in the center \**P* =0.0126; n = 7). (I-J) The number of entries to (L) and time spent in the center (M) by the hM4Di-expressing group during the Light-on and Light-off phases on the OFT (paired t-test; n = 10). (K) Schematic of the EPM paradigm. (L-M) The number of entries to (L) and time spent in the open arms (M) during the EPM test by the DISC1 control group during the Light-on and Light-off phases on the (paired t-test, number of entries \**P* = 0.0179, time spent in the open arms \*\**P* = 0.0031; n = 7). (N-O) The number of entries to (N) and time spent in the open arms (O) of the DISC1 experiment group in EPM (paired t-test; n = 7).

## Discussion

Epidemiological evidence has shown that many different mental illnesses share the same genetic risk (Gottesman et al., 2010; Lichtenstein et al., 2009) and share similar phenotypes (Hill et al., 2008; Ivleva et al., 2010), thus blurring the diagnostic and genetic boundaries of these illnesses (Porteous et al., 2011). There is still a lack of understanding regarding the neural basis of abnormal risk-avoidance behavior, a phenotype shared by multiple mental illnesses. In our study, we used the *DISC1-N^TM^* mouse model, which has an impairment in risk avoidance, to study the underlying neuronal mechanism. We found abnormal astrocytes-neurons communication in the BLA was responsible for the impaired risk avoidance observed in *DISC1-N^TM^* mice. Optogenetic stimulation of BLA ChR2-expressing astrocytes induced action potentials in WFS1 neurons and restored neuronal excitability, thereby rescuing the risk-avoidance impairment in *DISC1-N^TM^* mice. In this process, astrocytes regulate WFS1 neurons through NMDA receptors, and WFS1 neurons are required for astrocytes in the BLA to rescue abnormal risk avoidance.

Many paradigms have been used to assess approach-avoidance conflict in rodents, mainly including punishment-induced conflict paradigms (e.g., Vogel conflict tests), exploratory conflict paradigms (e.g., the EPM), and social interaction conflict paradigms (e.g., the social interaction test) (Kirlic et al., 2017b). The three paradigms we used to test risk-avoidance behavior in *DISC1-N^TM^* mice are exploratory conflict paradigms (OFT, EPM and a light-dark box). In these paradigms, rodents are faced with the desire to explore novel environments and a concurrent desire to avoid potential unknown threats (Kirlic et al., 2017a). Rodents have a natural aversion to open, elevated or well-lit areas as it makes them easier to be spotted by predators, more likely to fall and become injured and makes potential escape more difficult. However, exploring new areas can bring benefits, such as increasing chances of successful foraging or finding potential mates. Thus, an approach-avoidance conflict arises. Risk-avoidance behavior is assessed by monitoring frequency of visits to the risky area and the duration of time spent there (the central area in the OFT, the open-arms of the EPM and the light compartment in the LDB). While these exploratory conflict paradigms were developed to assess risk-preference behavior, increased avoidance to the risky areas is commonly thought to be associated with an anxiety-related response (Calhoon and Tye, 2015; Walf and Frye, 2007). In our study, *DISC1-N^TM^* mice showed reduced avoidance to the risky areas compared to control mice, whereas responses influenced by anxiety are highlighted by increased risk avoidance. Previous studies of motivation and behavior have established that increased novelty-seeking behavior in mice can enhance exploration of new objects or environments (Kliethermes and Crabbe, 2006; Pogorelov et al., 2005), resulting in similar reduced risk-avoidance performance in the conflict-avoidance paradigms. High novelty-seeking behavior is a psychopathological indicator for mental disorders such as drug abuse or bipolar disorder (Abreu-Villaça et al., 2006; Bardo et al., 1996; Minassian et al., 2011). To determine whether *DISC1-N^TM^* mice had high novelty-seeking behavior, we used a novel object recognition test of exploration behavior (Figure 1L-N) and found that *DISC1-N^TM^* mice did not have elevated exploratory behavior to novel objects, thereby excluding the effect of high novelty-seeking on risk-avoidance performance.

In our study, *DISC1-N^TM^* mice had low risk-avoidance performance (Figure 1A-L). However, normal risk avoidance or high risk avoidance has also been observed in other DISC1 models, for example, in the dominant-negative DISC1 transgenic model (DN-DISC1) (Hikida et al., 2007) or the DISC1 overexpressed model (Wang et al., 2019). In studies using DISC1 mutant models, studies of behavioral exploration were mostly associated with mental-illnesses-related phenotypes (or abnormal performance related to mental illnesses), such as approach-avoidance behavior, anhedonia, social behavior, startle response, and learning and memory deficits (Gómez-Sintes et al., 2014; Hikida et al., 2007; Kaminitz et al., 2014; Lipina et al., 2010; Wang et al., 2019). Different DISC1 models had different performance levels in the same behavioral paradigm, which we suspect was due to the complexity of the DISC1 protein. As an important hub protein, DISC1 interacts with many molecules and participates in multiple physiological activities (Niwa et al., 2016), including neurogenesis (Kim et al., 2009; Ye et al., 2017), synaptic plasticity (Tropea et al., 2018) and neuronal signaling (Kim et al., 2009; Lipina et al., 2013). Various DISC1 protein functions involving different interacting proteins may be affected in different transgenic models, resulting in different behavioral phenotypes. The complex functions of the DISC1 protein and the diversity of phenotypes in DISC1 transgenic models also reflect the complexity of pathogenesis of mental illnesses. The DISC1 model we used in this study was the *DISC1-N* terminal fragment transgenic mouse model. The N-terminal of the DISC1 protein is required for nuclear targeting of the DISC1 protein (Millar et al., 2005) and can interact with glycogen synthase kinase-3 (GSK-3, a serine/threonine protein kinase) (Mao et al., 2009). A pertinent outstanding question regarding the molecular mechanisms of the *DISC1* gene in abnormal risk-avoidance behavior is to clarify whether the abnormal interaction between astrocytes and neurons is due to the overexpression of the DISC1-N terminals or is due to the disfunction of intact DISC1 protein.

The four molecular for neuronal subtypes markers we chose (Figure 5A, B, S7A-D) have been found to participate in the pathology of mental illnesses. Calb2 neurons play a role in reward deficits in mood disorders (Russo and Nestler, 2013) and the abnormal neuronal function can lead to changes in substance abuse behavior (Konig et al., 2020). Abnormalities in RNA editing of *Htr2C* have been detected in suicide victims that suffer from depression (Iwamoto and Kato, 2003) and this gene is a hopeful candidate genes for investigating the mechanisms underlying mental illness (Iwamoto et al., 2009). MAP4K3, a serine/threonine kinase, is not only an important modulator of autophagy (Hsu et al., 2018; Lam et al., 2009), but is also involved in the response to environmental stress (Diener et al., 1997). Autophagy is associated with many mental illnesses, such as schizophrenia (Merenlender-Wagner et al., 2015). WFS1 protein is an endoplasmic reticulum (ER) membrane-embedded protein and has physiological functions in protein biosynthesis/processing, modulating ER calcium homeostasis and in the regulation of ER stress (Fonseca et al., 2005; Takei et al., 2006; Zatyka et al., 2008). Mutations in the *Wfs1* gene are associated with wolfram syndrome, an autosomal recessive disorder, which can also be sporadic, that affects many systems in the body including the CNS (Rigoli et al., 2011). About 60% of patients with Wolfram syndrome have symptoms associated with psychiatric disorders, such as severe depression, psychosis, and aggression (Swift et al., 1990), and asymptomatic mutation carriers exhibit higher rates of depression and attempted suicide (Swift and Gorman Swift, 2000). *Wfs1* gene mutations or functional changes have also been observed in mental illness, such as post-traumatic stress disorder (PTSD) (Kesner et al., 2009), depression and bipolar disorder (Koido et al., 2005). We have found that the high-response neurons recorded by patch clamp in the BLA were WFS1 neurons (Figure 5B) and that the excitability of WFS1 neurons was lower in *DISC1-N^TM^* mice thank in control mice (Figure 4H, I). However, we still do not know whether aberrant DISC1-N protein expression in *DISC1-N^TM^* mice is associated with reduced WFS1 neuronal excitability.

There has been an increase in studies reporting astrocyte participation in the function of neural circuits and behavioral outputs. In these studies, astrocytes were found to be heterogeneous. A diversity of morphology and gene expression in astrocytes has been confirmed both *in vitro* and *in vivo* (Bachoo et al., 2004; Matyash and Kettenmann, 2010; Raff et al., 1983; Rusnakova et al., 2013; Yong et al., 1990). At the functional level, a study showed that adult rat neural stem cells can differentiate into neurons when co-cultured with hippocampal astrocytes but not when co-cultured with spinal-cord-derived astrocytes (Song et al., 2002). Astrocytes can recognize different neuronal subtypes, and regulate specific types of neurons within a brain region (Eachus et al., 2017; Tan et al., 2017). For example, selective stimulation ChR2-expressing astrocytes in the CA1 area specifically increases the firing frequency of CCK but not parvalbumin interneurons (Tan et al., 2017). In addition to regulating different neurons, astrocytes are themselves regulated by specific neurons and activated by specific animal behaviors (Nimmerjahn et al., 2009). One study focused on the mouse dorsal striatum showed that different astrocyte subsets selectively respond to the activity of medium spiny neuron (MSN) (Martín et al., 2015). Astrocytes have inter- and intra-regional distinctions (Haim and Rowitch, 2017). Therefore, heterogeneity of functions must be considered when studying astrocytes, especially in investigations focused on a specific brain region or neural circuit. We have found that BLA astrocytes exhibit cell-type-specific modulation of nearby neuronal populations (that is, high-response neurons and low-response neurons). We confirmed that high-response neurons are WFS1 neurons and are involved in risk-avoidance behavior, but the molecular markers and functions of low-response neurons still require further exploration. We speculate that photostimulation-induced ChR2-expression of astrocytes in the BLA may lead to the release of more than one gliotransmitters which act on nearby neurons, binding to low-response neurons and causing slight depolarization of the cell membrane.

Optogenetics is a combination of genetic manipulation techniques and optical methods which can specifically activate or inhibit neurons (Deisseroth, 2011). This technique has been widely used in neuroscience research since it was first described in 2005 (Boyden et al., 2005). ChR2 is one of the most commonly used light-sensitive proteins in optogenetics. ChR2 is a cation channel that is excited by light around 473 nm, and has high permeability to monovalent cations (Na^+^, K^+^ and protons) (Nagel et al., 2003). With the widespread application of optogenetics, light-sensitive proteins like ChR2 have also been expressed on astrocytes in functional studies. Photostimulation of ChR2-expressing astrocytes can potently induce sleep during the active phase of the sleep-wake cycle (Pelluru et al., 2016). In the primary visual cortex, photostimulation of ChR2-expressing astrocytes can enhance excitatory and inhibitory synaptic transmission by activating metabotropic glutamate receptors (Perea et al., 2014). In a functional magnetic resonance imaging (fMRI) study, photostimulation of ChR2-expressing astrocytes alone could elicit blood oxygenation level-dependent (BOLD) signaling responses (Takata et al., 2018). Both *in vitro* and *in vivo* evidence has suggested that ChR2 selectively stimulates astrocytes by inducing an increase in intracellular Ca^2+^ concentration, which in turn modulates neuronal function and even participates in behavior (Chen et al., 2013; Gourine et al., 2010). But the exact mechanism of how ChR2 enhances astrocyte functions is unclear. Studies have suggested that the increase of Ca^2+^ concentration induced by ChR2 in astrocytes mainly comes from the release of intracellular stored Ca^2+^, and the ATP released by photostimulated astrocytes can bind to P2Y receptors via autocrine signalling, which in turn increases the concentration of intracellular Ca^2+^ (Figueiredo et al., 2014). However, it has also been found that the ChR2-induced increase in intracellular Ca^2+^ concentration in astrocytes is predominantly mediated by the Na^+^-Ca^2+^ exchangers caused by the influx of Na^+^ through ChR2 channels (Yang et al., 2015). In the absence of extracellular Ca^2+^, photostimulation failed to induce intracellular Ca^2+^ concentration increase in ChR2-expressing astrocytes (Yang et al., 2015). Although the mechanism by which ChR2 leads to elevated astrocyte function remains unclear, it is well established that ChR2 initiates biologically relevant signaling events in astrocytes and can be used as a tool to explore astrocyte function. We used a virus vector to express ChR2 on astrocytes and found photostimulation astrocytes in the BLA can rescue abnormal risk-avoidance behavior in *DISC1-N^TM^* mice.

To explore the mechanism underlying astrocyte-neuron interactions, we used the patch clamp technique to record adjacent neurons while optogenetically stimulating ChR2-expressing astrocytes in the BLA. We found that light-stimulated astrocytes modulate neuronal activity through the glycine binding site of NMDARs. NMDARs are a key player in excitatory transmission and synaptic plasticity in the CNS (Hunt and Castillo, 2012; Paoletti et al., 2013). When NMDARs bind glutamate to its glutamate binding site and a co-agonist such as D-serine to its glycine binding site, the receptors become activated and cations can pass through the cell membrane, leading to depolarization of the membrane potential (Furukawa et al., 2005). In organisms, D-serine is an important endogenous NMDAR ligand, and it is present in the brain in quantities higher even than many essential amino acids (Hashimoto et al., 1992; Hashimoto et al., 1993). Astrocytes are one of the main sources of D-serine in the CNS (Papouin et al., 2017b). Astrocytes contain serine racemase, which can convert L-serine to D-serine and release D-serine into the synaptic cleft, affecting the electrophysiological activity of neurons (Miller, 2004; Panatier et al., 2006; Wolosker et al., 2002). Some studies also propose that astrocytes synthesize L-serine which shuttles to neurons to fuel the neuronal synthesis of D-serine (serine shuttle model) (Wolosker, 2011; Wolosker et al., 2016), which is then released by the neurons and can be taken up by astrocytes for storage and activity-dependent release (Wolosker, 2011). Studies have found that D-serine released by astrocytes can affect long-term potentiation (Henneberger et al., 2010; Kang et al., 2013) and affect behaviors related to mental illnesses (for example, depression (Otte et al., 2013) and anxiety-related behavior (Otte et al., 2013)). We speculate that astrocytes may influence the activity of WFS1 neurons using D-serine.

In summary, our work indicates that astrocytes in the BLA may be an effective target for the treatment of abnormal risk-avoidance behavior. Moreover, our results show that neurons have differential effects on nearby astrocytes, suggesting that BLA astrocytes are heterogeneous and may be involved in different behaviors through different neuronal subtypes. This emphasizes the complexity of astrocyte function in modulating neural circuits.

## Acknowledgements

We thank Dr. Weidong Li for *DISC1-N^TM^* transgenic mice. This study was supported by the National Natural Science Foundation of China (31671116 JT, 31761163005 JT), the Scientific Instrument Developing Project of the Chinese Academy of Sciences (YJKYYQ20200109115405930 JT), the China Postdoctoral Science Foundation (2022M712188 XY. Z), the Guangdong Provincial Key S&T Program (2018B030336001 JT), Science and Technology Program of Guangzhou (202007030001 JT), the Key Basic Research Program of Shenzhen Science and Technology Innovation Commission (JCYJ20200109115405930 JT; JCYJ20200109150717745 XY).

## Author contributions

J.T. conceived the study. X-Y.Z., Q. X., S.C., X-R.X. and Y-C.H. performed experiments. X-Y. Z., S.C. and Y-W. C. analyzed data. F. Y., Y. C. and L-P. W. provided suggestions and comments on the manuscript. X-Y.Z. and J.T. wrote the manuscript. J.T. supervised the project.

## Materials and Methods

### Animals

We used the following mouse lines: inducible *DISC1-N* terminal fragment transgenic mice (*DISC1-N^TM^*, male, generated by Prof. Weidong Li’s lab, Shanghai Jiao Tong University) and C57BL/6J mice (male, Guangdong Medical Laboratory Animal Center, Guangzhou, China).

Mice were given free access to food pellets and water and were maintained on a 12 h/12 h light/dark cycle (lights on at 8:00 a.m.). All experiments were approved by the Shenzhen Institutes of Advanced Technology, Chinese Academy of Sciences Research Committee, and all experimental procedures involving animals were carried out in strict accordance with the Research Committee’s animal use guidelines. Surgeries were performed under full anesthesia and every effort was made to minimize animal suffering.

### Behavioral tests

All behavioral tests were performed blind to mice genotype. Groups of mice were age-matched (2–4 months) and, prior to behavioral assays, were handled for 5 min per day for three days to reduce stress introduced by experimenter contact. Tamoxifen (Sigma-Aldrich, #T5648) was administered to all mice (intraperitoneal injection, 0.05mg/g, drug concentration: 0.005 g/ml in corn oil) six hours before behavioral testing unless indicated otherwise. In an oil control group, mice were injected with the same volume of corn oil (Sigma, C8267-500ML) with no tamoxifen. Behavioral tests using CNO (MedChemExpress, #HY-17366) were conducted in a 60-min window that began 30 min after CNO administration (intraperitoneal injection, 1 mg/kg). The ambient light was around 200 lux during all behavioral tests.

#### 1) Open field test (OFT)

The open field consisted of an unmarked box made of white acrylic board with a length, width and height of 48 cm. At the beginning of the experiment, mice were placed in a fixed corner of the box, allowing free exploration of the space for 5 min (exploration phase). The subsequent photostimulation phase also lasted for 5 min and there was no interval between the two phases. The behavioral tests were recorded using a video camera directly above and videos were analyzed using ANY-maze software (Stoelting, U.S.A).

#### 2) Elevated plus maze (EPM)

A plastic elevated plus maze was used, consisting of a central platform (5×5 cm) with two white open arms (25×5 cm) and two white closed arms (25×5 cm, with 17-cm-high surrounding walls) extending from it to form a plus shape. The maze was elevated 58 cm above the floor. Mice were placed in the central area of the EPM with their heads facing an open arm and were allowed to freely explore for 3 min. The subsequent photostimulation phase also lasted for 3 min and there was no interval between the two phases. The behavioral tests were recorded using a video camera directly above and videos were analyzed using ANY-maze software.

#### 3) Light-dark box (LDB)

A light-dark box consisted of a black acrylic box and a white acrylic box of the same size (length 20cm, width 20cm, height 14cm). The white box had no lid whereas a lid was provided above the black box and closed during the experiment. There was a removable door (7×7cm) separating the two boxes. At the beginning of the test, mice were placed in the dark box and the door was removed after 6 s so that mice could explore both boxes. The duration of the experiment was 5 min.

#### 4) Novel object recognition task (NOR)

The novel object recognition task was carried out in an unmarked open box made of white acrylic sheet material (48 × 48 × 48 cm). The objects (toys) were similar in size, around 5.5 cm × 5.5 cm × 5.5 cm (Rubik’s cubes, plastic dolls, and transparent plastic boxes with colored clips). Each experiment lasted 4 consecutive days, and each session lasted 10 min. On the first day, each mouse was placed in a corner (used as a starting position in all sessions/mice) and was free to explore the empty box. On the second and third days, mice were presented with objects during an object familiarization stage; two objects were placed in the corners of the box (diagonally opposite corners, 5 cm away from the two walls) and the mice were free to explore the box or toys. On the fourth day, one familiar object was replaced by a novel object and the exploration times for each object was recorded. If the distance between the mouse’s nose and the toy was less than 2 cm, the mouse was considered to be exploring the toy.

#### 5) Looming test

We used an automatic behavioral detection system established in our laboratory to detect looming behavior in mice (Yang et al., 2020). The box for testing the behavioral performance of mice consisted of an open cylinder (diameter: 50 cm, height: 30 cm) adjacent to rectangular box (length: 50 cm, width: 10 cm, height: 30 cm). Visual stimuli (black) were presented on a 42-inch. LCD monitor (refresh rate 60 Hz, AOC) displaying a gray background, positioned 46 cm above the arena floor. The arena was built around an infrared touchscreen frame, which was positioned approximately at the height of a mouse body and recorded the coordinates of the locations visited by each mouse. The recording software was programmed in C++ using Visual Studio 2015.

In the beginning of the experiment, mice were put in the rectangular box and were free to explore the arena for 5 min. When mice entered a circular trigger area (diameter 25 cm), the looming stimulus was automatically triggered (a rapidly expanding black disc on screen, expanding from a visual angle of 2° to 40° in 250 ms, i.e., an expanding speed of 152°/s). The expanding disc stimulus was repeated for 15 times in quick succession. Each experimental mouse experienced 5 sets of looming stimuli within 30 minutes, and an interval of ≥3 minutes between each stimulus. We selected the data from the first three looming stimulations from each mouse for statistical analysis.

#### 6) Sucrose preference test

The sugar water preference test took place inside a standard mouse cage. Two identical drinking bottles were placed on the cage and mice could freely choose to drink from either. Five days before the start of the experiment, mice were given a 48-hour supply of drinking water using two drinking bottles, and then another 48-hour supply of two bottles containing 2% sucrose water. The day before the experiment started, mice were deprived of water for 24 hours. On the day of the experiment, mice were placed in a single cage and two drinking bottles were available, one with drinking water and the other with 2% sugar water. Before the start of the experiment, all the water bottles were marked and weighed. At 3 hours into the session, the positions of the drinking bottles were exchanged to prevent the mice from developing positional preference for the bottles. After the 6-hour timepoint, all bottles were weighed again, and the liquid consumed from each bottle was calculated.

### Stereotactic virus injection and optogenetic manipulation

Adeno-associated viruses (AAVs) carrying GFAP promoter (AAV-GFAP-ChR2-mCherry, titers 3×10^12^ particles per ml, AAV-GFAP-ChR2-eYFP, titers 3×10^12^ particles per ml, AAV-GFAP-hM4Di-mCherry, titers 3×10^12^ particles per ml, AAV-GFAP-mCherry, titers 3×10^12^ particles per ml) were packaged in our laboratory. AAV-Wfs1-hM4Di-mCherry and AAV-Wfs1-mCherry were packed by BrainVTA (China). Mice were deeply anesthetized with 1% sodium pentobarbital (Sigma-Aldrich, #P3761, 10 ml/kg body weight, i.p.) and placed in a stereotaxic instrument (RWD Life Science Inc., Shenzhen, China) and head fixed. A microinjector pump (UMP3/Micro4, USA) was used to pump a microliter syringe (10 μl, Hamilton) to inject virus to the target region at a speed of 80 nl/min. The injection needle was left in place for 10 min at the end of injection to avoid reflux of the viral solution. A volume of 200 nl of AAV-GFAP-ChR2-mCherry or AAV-Wfs1-mCherry was unilateral injected into the BLA (AP +3.00 mm, ML -1.22 mm, DV −5.00 mm) for optogenetic and electrophysiological experiments. Similarly, 200 nl of AAV-Wfs1-hM4Di-mCherry or AAV-Wfs1-mCherry was bilateral injected into the BLA (AP +3.00 mm, ML -1.22 mm, DV −5.00 mm) for designer receptors exclusively activated by a designer drug (DREADD) experiment. For optogenetic experiments, two weeks after virus injection, mice were implanted with a 200 μm unilateral fiber optic cannula (AP +3.00 mm, ML -1.22 mm, DV −4.80 mm) secured to the skull with denture base material (SND, China) and dental base acrylic resin powder (Feiying, China). Mice were then given one week to recover before behavioral experiments began. In the optogenetic stimulation behavior test, blue light stimulation (wavelength: 470 nm, frequency: 20 Hz, width: 25 ms, power: 5-8 mW) was turned on and directed through the optic cannula at the BLA during the “light on” stage. A control (mCherry) group underwent the same procedure and received the same intensity of laser stimulation.

### Immunohistochemistry

Mice received an overdose of 1% sodium pentobarbital (15 ml/kg body weight, i.p.) and were transcardially perfused with phosphate-buffered saline (PBS), followed by 4% paraformaldehyde (PFA; Aladdin, #C104188) in PBS. Brains were removed and submerged in 4% PFA at 4 °C overnight to post-fix, and then transferred to 20% sucrose for one day and then 30% sucrose for 2 days. We used O.C.T. compound (Tissue-Tek^®^, O.C.T optimal cutting temperature) to quickly embed brain tissues before cutting frozen sections. Coronal brains sections (30-μm thickness) were obtained using a cryostat microtome (Lecia CM1950, Germany). Brain sections were washed with PBS (3 min, room temperature) 3 times to wash out the O.C.T. Then, brain sections were put into blocking solution (0.3% Triton X-100 and 10% normal goat serum, NGS in PBS, 1 hour at room temperature). Brain sections were then incubated overnight in primary antiserum (rabbit anti-S100, 1:300, Abcam; mouse anti-NeuN, 1:300, Abcam; rabbit anti-WFS1, 1:300, proteintech) diluted in PBS with 3% NGS and 0.1% Triton X-100. The following day, the sections were incubated in secondary antibodies at room temperature for 1 hour. The secondary antibodies used were Alexa Fluor^®^ 488, 594 or 647-conjugated goat anti-rabbit or anti-mouse IgG antibodies (1:300, Invitrogen, CA, USA) at room temperature for 1 hour. Then, brain sections were mounted and covered slipped with anti-fade reagent with DAPI (ProLong Gold Antifade Reagent with DAPI, life technologies). Brain sections were then photographed and analyzed using an Olympus slide scanner VS120-S6-W or Leica TCS SP5 laser scanning confocal microscope. Definitions of brain regions were based on The Mouse Brain in Stereotaxic Coordinates (Franklin and Paxinos, 1997).

### Patch-clamp electrophysiology

For patch clamp recording, all drugs used were from Sigma-Aldrich unless indicated otherwise. Coronal slices (270 μm) containing BLA (bregma -0.7 to – 1.9 mm) were prepared using standard procedures. Brains were quickly removed and chilled in ice-cold modified artificial cerebrospinal fluid (ACSF) containing (in mM): 110 Choline Chloride, 2.5 KCl, 1.3 NaH_2_PO_4_, 25 NaHCO_3_, 1.3 Na-Ascorbate, 0.6 Na-Pyruvate, 10 Glucose, 2 CaCl_2_, and 1.3 MgCl_2_. Then the BLA slices were cut in ice-cold modified ACSF using a Leica vibroslicer (VT-1200S). Slices were allowed to recover for 30 min in a storage chamber containing regular ACSF at 32–34 °C (in mM): 125 NaCl, 2.5 KCl, 1.3 NaH_2_PO_4_, 25NaHCO_3_, 1.3 Na-Ascorbate, 0.6 Na-Pyruvate, 10 Glucose, 2 CaCl_2_, 1.3 MgCl_2_ (pH 7.3∼7.4 when saturated with 95% O_2_/5% CO_2_), and thereafter kept at room temperature, until placed in the recording chamber. The osmolarity of all the solutions was 300∼320 mOsm/kg. For all electrophysiological experiments, slices were viewed using infrared optics under an upright microscope (Eclipse FN1, Nikon Instruments). The recording chamber was continuously perfused with oxygenated ACSF (2 ml/min) at room temperature. Pipettes were pulled using a micropipette puller (Sutter P-2000 Micropipette Puller) with a resistance of approximately 5-10 MΩ. Recordings were made with electrodes filled with intracellular solution (in mM): 130 potassium gluconate, 1 EGTA, 10 NaCl, 10 HEPES, 2 MgCl_2_, 0.133 CaCl_2_, 3.5 Mg-ATP, and 1 Na-GTP.

Action potential firing frequency was analyzed in current-clamp mode in response to a 2 s depolarizing current step. All recordings were conducted with a MultiClamp700B amplifier (Molecular Devices). Currents were low-pass filtered at 2 kHz and digitized at 20 kHz using an Axon Digidata 1440A data acquisition system and pClamp 10 software (both from Molecular Devices). Series resistance (Rs) was approximately 10–30 MΩ and regularly monitored throughout the recordings. Data were discarded if the series resistance (Rs) changed by >30% over the course of data acquisition.

### Real-time fluorescence quantitative PCR (qPCR)

#### 1) qPCR of BLA brain tissue

Six hours after intraperitoneal injection of tamoxifen, mice were anesthetized using isoflurane (3%) and the heads were decapitated to obtain the brain tissue, which were sliced with a vibroslicer (Leica, VT-1200S) to obtain brain slices containing the BLA region. Brain slices were obtained in the same way as in patch clamp. Slices were placed on ice and immersed in ACSF saturated with 95% O_2_-5% CO_2_, and both sides of the BLA were separated with forceps and surgical blades (RWD Life Science, Shenzhen, China) under a stereo microscope (Zeiss).

After enriching the BLA tissue of three mice, we used an RNA extraction kit (Transgen, TransZol Up Plus RNA Kit) and qPCR kit (Transgen, TransStart® Green qPCR SuperMix) for RNA extraction and reverse transcription. The LightCycler 480 (Roche) was used to for real-time PCR.

#### 2) qPCR for a single cell

After the electrophysiological characteristics of neurons were recorded in patch clamp recordings, negative pressure was then applied to the neurons through the glass electrodes, slowly lifting the glass electrodes away from the brain slice. Target neurons can be “sucked” from the brain slice in this way. Following careful removal of the glass electrode with a single target neuron at the tip, we placed it into the pre-cooled lysis solution in a PE tube. The glass electrode directly contacted the bottom of the tube and the tip of the glass electrode was broken in the tube. Next, the extracted cells were reverse-transcribed and amplified according to the standard steps in the reverse transcription kit (single cell-to-CT kit, ambion). The sample was placed on the LightCycler 480 (Roche) and the protocol instructions were followed for the reverse transcription kit to obtain the qPCR results.

We adopted a relative quantitative method to analyze the results of the qPCR experiments between WT and *DISC1-N^TM^* mice. The δCt values were used to compare the difference in gene expression between the test samples (Ct stands for the cycle threshold, “target” represents the target gene, “reference” represents the internal reference gene).

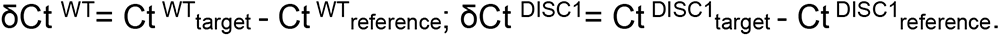

The qPCR of BLA brain tissue used β-actin as the internal reference gene, and the qPCR of a single cell used RNA 18s as the internal reference gene.

### Single-nucleus RNA sequencing (snRNA-seq)

Brain slices were obtained in the same way as in qPCR of BLA brain tissue and the acquired BLA tissue was stored frozen on dry ice pending snRNA-seq. To obtain a sufficient number of nuclei, we enriched bilateral BLA brain tissue from 5 mice for each group. Frozen brain tissue was placed directly into a pre-chilled Dounce homogenizer containing ice-cold homogenization buffer (0.25M sucrose, 25mM KCl, 5mM MgCl_2_, 20mM tricine-KOH [pH 7.8], 1mM dithiothreitol, 0.15mM spermine, 0.5mM spermidine, protease inhibitors, 5μg/mL actinomycin, 0.32% Nonidet P-40, and 0.04% bovine serum albumin). After 25 beats with a pestle, the homogenate was mixed 1:1 in OptiPrep and centrifuged at 10,000× g for 20 min at 4 °C. Nuclei were collected at the bottom of the centrifuge tube, washed once to remove OptiPrep, and resuspended in Dulbecco’s modified eagle medium/F12 supplemented with 10% fetal bovine serum. We diluted the nuclei to 400 nuclei per microliter using a hemocytometer. All buffers and gradients used for nuclear extraction contained RNase inhibitor (Promega) at 60 U/mL.

We generated snRNA-seq libraries using the Chromium Single Cell 3’ Library Kit v3 (1000078; 10x Genomics) according to the manufacturer’s instructions. We used 40 μL of the diluted nucleus suspension mixed with the reverse transcription reagent mix and loaded the samples into the chip for single-cell nuclei encapsulation. The encapsulated nuclei were incubated on a PCR thermocycler to allow reverse transcription of RNA to generate barcoded cDNA. We used these cDNAs for library construction; the final library concentration was determined by Qubit (Thermo Fisher Scientific), and the fragment length was determined using a Fragment Analyzer (Advanced Analytical Technologies). We subjected libraries to paired-end sequencing on the NovaSeq 6000 System according to the manufacturer’s instructions (Novogene) and obtained at least 200 GB of raw data per library.

## Notes

### Competing Interest Statement

The authors have declared no competing interest.

## Reference

(2000). Cross-national comparisons of the prevalences and correlates of mental disorders. WHO International Consortium in Psychiatric Epidemiology. Bull World Health Organ 78, 413–426.

Abreu-Villaça, Y., Queiroz-Gomes, F.d.E., Dal Monte, A.P., Filgueiras, C.C., and Manhães, A.C. (2006). Individual differences in novelty-seeking behavior but not in anxiety response to a new environment can predict nicotine consumption in adolescent C57BL/6 mice. Behavioural Brain Research 167, 175–182.

Allen, N.B., and Badcock, P.B. (2003). The social risk hypothesis of depressed mood: evolutionary, psychosocial, and neurobiological perspectives. Psychological bulletin 129, 887.

Bachoo, R.M., Kim, R.S., Ligon, K.L., Maher, E.A., Brennan, C., Billings, N., Chan, S., Li, C., Rowitch, D.H., Wong, W.H., et al. (2004). Molecular diversity of astrocytes with implications for neurological disorders. 101, 8384–8389.

Bardo, M.T., Donohew, R.L., and Harrington, N.G. (1996). Psychobiology of novelty seeking and drug seeking behavior. Behavioural brain research 77, 23–43.

Blomeley, C., Garau, C., and Burdakov, D. (2018). Accumbal D2 cells orchestrate innate risk-avoidance according to orexin signals. Nature Neuroscience 21, 29–32.

Boyden, E.S., Zhang, F., Bamberg, E., Nagel, G., and Deisseroth, K. (2005). Millisecond-timescale, genetically targeted optical control of neural activity. Nature Neuroscience 8, 1263–1268.

Calhoon, G.G., and Tye, K.M. (2015). Resolving the neural circuits of anxiety. Nature Neuroscience 18, 1394–1404.

Chen, J., Tan, Z., Zeng, L., Zhang, X., He, Y., Gao, W., Wu, X., Li, Y., Bu, B., Wang, W., et al. (2013). Heterosynaptic long-term depression mediated by ATP released from astrocytes. Glia 61, 178–191.

Deisseroth, K. (2011). Optogenetics. Nature Methods 8, 26–29.

Diehl, M.M., Iravedra-Garcia, J.M., Morán-Sierra, J., Rojas-Bowe, G., Gonzalez-Diaz, F.N., Valentín-Valentín, V.P., and Quirk, G.J. (2020). Divergent projections of the prelimbic cortex bidirectionally regulate active avoidance. Elife 9, e59281.

Diener, K., Wang, X.S., Chen, C., Meyer, C.F., Keesler, G., Zukowski, M., Tan, T.H., and Yao, Z. (1997). Activation of the c-Jun N-terminal kinase pathway by a novel protein kinase related to human germinal center kinase. Proceedings of the National Academy of Sciences of the United States of America 94, 9687–9692.

Duff, B.J., Macritchie, K.A.N., Moorhead, T.W.J., Lawrie, S.M., and Blackwood, D.H.R. (2013). Human brain imaging studies of DISC1 in schizophrenia, bipolar disorder and depression: A systematic review. Schizophrenia Research 147, 1–13.

Eachus, H., Bright, C., Cunliffe, V.T., Placzek, M., Wood, J.D., and Watt, P.J. (2017). Disrupted-in-Schizophrenia-1 is essential for normal hypothalamic-pituitary-interrenal (HPI) axis function. Human molecular genetics 26, 1992–2005.

Eaton, W.W., Bienvenu, O.J., and Miloyan, B. (2018). Specific phobias. The lancet Psychiatry 5, 678–686.

Endrass, T., Schuermann, B., Roepke, S., Kessler-Scheil, S., and Kathmann, N. (2016). Reduced risk avoidance and altered neural correlates of feedback processing in patients with borderline personality disorder. Psychiatry Research 243, 14–22.

Figueiredo, M., Lane, S., Stout, R.F., Liu, B., Parpura, V., Teschemacher, A.G., and Kasparov, S. (2014). Comparative analysis of optogenetic actuators in cultured astrocytes. Cell Calcium 56, 208–214.

Fonseca, S.G., Fukuma, M., Lipson, K.L., Nguyen, L.X., Allen, J.R., Oka, Y., and Urano, F. (2005). WFS1 is a novel component of the unfolded protein response and maintains homeostasis of the endoplasmic reticulum in pancreatic beta-cells. The Journal of biological chemistry 280, 39609–39615.

Furukawa, H., Singh, S.K., Mancusso, R., and Gouaux, E. (2005). Subunit arrangement and function in NMDA receptors. Nature 438, 185–192.

Gómez-Sintes, R., Kvajo, M., Gogos, J.A., and Lucas, J.J. (2014). Mice with a naturally occurring DISC1 mutation display a broad spectrum of behaviors associated to psychiatric disorders. Front Behav Neurosci 8, 253.

Gottesman, I.I., Laursen, T.M., Bertelsen, A., and Mortensen, P.B. (2010). Severe Mental Disorders in Offspring With 2 Psychiatrically Ill Parents. Archives of General Psychiatry 67, 252–257.

Gourine, A.V., Kasymov, V., Marina, N., Tang, F., Figueiredo, M.F., Lane, S., Teschemacher, A.G., Spyer, K.M., Deisseroth, K., and Kasparov, S. (2010). Astrocytes control breathing through pH-dependent release of ATP. Science (New York, NY) 329, 571–575.

Grant, S., Contoreggi, C., and London, E.D. (2000). Drug abusers show impaired performance in a laboratory test of decision making. Neuropsychologia 38, 1180–1187.

Haim, L.B., and Rowitch, D.H. (2017). Functional diversity of astrocytes in neural circuit regulation. Nature Reviews Neuroscience 18, 31–41.

Hashimoto, A., Nishikawa, T., Hayashi, T., Fujii, N., Harada, K., Oka, T., and Takahashi, K. (1992). The presence of free D-serine in rat brain. FEBS letters 296, 33–36.

Hashimoto, A., Nishikawa, T., Oka, T., and Takahashi, K. (1993). Endogenous D-serine in rat brain: N-methyl-D-aspartate receptor-related distribution and aging. Journal of neurochemistry 60, 783–786.

Hashimoto, R., Numakawa, T., Ohnishi, T., Kumamaru, E., Yagasaki, Y., Ishimoto, T., Mori, T., Nemoto, K., Adachi, N., Izumi, A., et al. (2006). Impact of the DISC1 Ser704Cys polymorphism on risk for major depression, brain morphology and ERK signaling. Human Molecular Genetics 15, 3024–3033.

Hennah, W., Thomson, P., McQuillin, A., Bass, N., Loukola, A., Anjorin, A., Blackwood, D., Curtis, D., Deary, I.J., Harris, S.E., et al. (2009). DISC1 association, heterogeneity and interplay in schizophrenia and bipolar disorder. Molecular Psychiatry 14, 865–873.

Henneberger, C., Papouin, T., Oliet, S.H., and Rusakov, D.A. (2010). Long-term potentiation depends on release of D-serine from astrocytes. Nature 463, 232–236.

Hikida, T., Jaaro-Peled, H., Seshadri, S., Oishi, K., Hookway, C., Kong, S., Wu, D., Xue, R., Andradé, M., Tankou, S., et al. (2007). Dominant-negative DISC1 transgenic mice display schizophrenia-associated phenotypes detected by measures translatable to humans. Proceedings of the National Academy of Sciences of the United States of America 104, 14501–14506.

Hill, S.K., Harris, M.S.H., Herbener, E.S., Pavuluri, M., and Sweeney, J.A. (2008). Neurocognitive Allied Phenotypes for Schizophrenia and Bipolar Disorder. Schizophrenia Bulletin 34, 743–759.

Hsu, C.L., Lee, E.X., Gordon, K.L., Paz, E.A., Shen, W.-C., Ohnishi, K., Meisenhelder, J., Hunter, T., and La Spada, A.R. (2018). MAP4K3 mediates amino acid-dependent regulation of autophagy via phosphorylation of TFEB. Nature Communications 9, 942.

Hubbard, D.T., Blanchard, D.C., Yang, M., Markham, C.M., Gervacio, A., Chun-I, L., and Blanchard, R.J. (2004). Development of defensive behavior and conditioning to cat odor in the rat. Physiology & Behavior 80, 525–530.

Hunt, D.L., and Castillo, P.E. (2012). Synaptic plasticity of NMDA receptors: mechanisms and functional implications. Current Opinion in Neurobiology 22, 496–508.

Ivleva, E.I., Morris, D.W., Moates, A.F., Suppes, T., Thaker, G.K., and Tamminga, C.A. (2010). Genetics and intermediate phenotypes of the schizophrenia—bipolar disorder boundary. Neuroscience & Biobehavioral Reviews 34, 897–921.

Iwamoto, K., Bundo, M., and Kato, T. (2009). Serotonin receptor 2C and mental disorders: Genetic, expression, and RNA editing studies. RNA Biology 6, 248–253.

Iwamoto, K., and Kato, T. (2003). RNA editing of serotonin 2C receptor in human postmortem brains of major mental disorders. Neuroscience Letters 346, 169–172.

Jha, M.K., Kim, J.-H., Song, G.J., Lee, W.-H., Lee, I.-K., Lee, H.-W., An, S.S.A., Kim, S., and Suk, K. (2018). Functional dissection of astrocyte-secreted proteins: Implications in brain health and diseases. Progress in neurobiology 162, 37–69.

Kaminitz, A., Barzilay, R., Segal, H., Taler, M., Offen, D., Gil-Ad, I., Mechoulam, R., and Weizman, A. (2014). Dominant negative DISC1 mutant mice display specific social behaviour deficits and aberration in BDNF and cannabinoid receptor expression. The world journal of biological psychiatry : the official journal of the World Federation of Societies of Biological Psychiatry 15, 76–82.

Kang, N., Peng, H., Yu, Y., Stanton, P.K., Guilarte, T.R., and Kang, J. (2013). Astrocytes release D-serine by a large vesicle. Neuroscience 240, 243–257.

Kesner, Y., Zohar, J., Merenlender, A., Gispan, I., Shalit, F., and Yadid, G. (2009). WFS1 gene as a putative biomarker for development of post-traumatic syndrome in an animal model. Molecular Psychiatry 14, 86–94.

Kim, J.Y., Duan, X., Liu, C.Y., Jang, M.-H., Guo, J.U., Pow-anpongkul, N., Kang, E., Song, H., and Ming, G.-l. (2009). DISC1 Regulates New Neuron Development in the Adult Brain via Modulation of AKT-mTOR Signaling through KIAA1212. Neuron 63, 761–773.

Kirlic, N., Young, J., and Aupperle, R.L. (2017a). Animal to human translational paradigms relevant for approach avoidance conflict decision making. Behav Res Ther 96, 14–29.

Kirlic, N., Young, J., and Aupperle, R.L. (2017b). Animal to human translational paradigms relevant for approach avoidance conflict decision making. Behaviour research and therapy 96, 14–29.

Kliethermes, C.L., and Crabbe, J.C. (2006). Genetic independence of mouse measures of some aspects of novelty seeking. Proceedings of the National Academy of Sciences of the United States of America 103, 5018–5023.

Koido, K., Kõks, S., Nikopensius, T., Maron, E., Altmäe, S., Heinaste, E., Vabrit, K., Tammekivi, V., Hallast, P., Kurg, A., et al. (2005). Polymorphisms in wolframin (WFS1) gene are possibly related to increased risk for mood disorders. International Journal of Neuropsychopharmacology 8, 235–244.

Konig, N., Bimpisidis, Z., Dumas, S., and Wallen-Mackenzie, A. (2020). Selective Knockout of the Vesicular Monoamine Transporter 2 (Vmat2) Gene in Calbindin2/Calretinin-Positive Neurons Results in Profound Changes in Behavior and Response to Drugs of Abuse. Front Behav Neurosci 14, 578443.

Lam, D., Dickens, D., Reid, E.B., Loh, S.H.Y., Moisoi, N., and Martins, L.M. (2009). MAP4K3 modulates cell death via the post-transcriptional regulation of BH3-only proteins. Proceedings of the National Academy of Sciences 106, 11978–11983.

Lichtenstein, P., Yip, B.H., Björk, C., Pawitan, Y., Cannon, T.D., Sullivan, P.F., and Hultman, C.M. (2009). Common genetic determinants of schizophrenia and bipolar disorder in Swedish families: a population-based study. The Lancet 373, 234–239.

Lipina, T.V., Fletcher, P.J., Lee, F.H., Wong, A.H.C., and Roder, J.C. (2013). Disrupted-In-Schizophrenia-1 Gln31Leu Polymorphism Results in Social Anhedonia Associated with Monoaminergic Imbalance and Reduction of CREB and β-arrestin-1,2 in the Nucleus Accumbens in a Mouse Model of Depression. Neuropsychopharmacology 38, 423–436.

Lipina, T.V., Niwa, M., Jaaro-Peled, H., Fletcher, P.J., Seeman, P., Sawa, A., and Roder, J.C. (2010). Enhanced dopamine function in DISC1-L100P mutant mice: implications for schizophrenia. Genes, brain, and behavior 9, 777–789.

Lorian, C.N., and Grisham, J.R. (2011). Clinical implications of risk aversion: An online study of risk-avoidance and treatment utilization in pathological anxiety. Journal of Anxiety Disorders 25, 840–848.

Lorian, C.N., and Grisham, J.R. (2012). The Safety Bias: Risk-Avoidance and Social Anxiety Pathology. Behaviour Change 27, 29–41.

Maner, J.K., and Schmidt, N.B. (2006). The Role of Risk Avoidance in Anxiety. Behavior Therapy 37, 181–189.

Mao, Y., Ge, X., Frank, C.L., Madison, J.M., Koehler, A.N., Doud, M.K., Tassa, C., Berry, E.M., Soda, T., Singh, K.K., et al. (2009). Disrupted in Schizophrenia 1 Regulates Neuronal Progenitor Proliferation via Modulation of GSK3β/β-Catenin Signaling. Cell 136, 1017–1031.

Martín, R., Bajo-Grañeras, R., Moratalla, R., Perea, G., and Araque, A. (2015). Circuit-specific signaling in astrocyte-neuron networks in basal ganglia pathways. 349, 730–734.

Matyash, V., and Kettenmann, H. (2010). Heterogeneity in astrocyte morphology and physiology. Brain Research Reviews 63, 2–10.

Merenlender-Wagner, A., Malishkevich, A., Shemer, Z., Udawela, M., Gibbons, A., Scarr, E., Dean, B., Levine, J., Agam, G., and Gozes, I. (2015). Autophagy has a key role in the pathophysiology of schizophrenia. Molecular Psychiatry 20, 126–132.

Millan, M.J. (2003). The neurobiology and control of anxious states. Progress in neurobiology 70, 83–244.

Millar, J.K., James, R., Christie, S., and Porteous, D.J. (2005). Disrupted In Schizophrenia 1 (DISC1): Subcellular targeting and induction of ring mitochondria. Molecular and Cellular Neuroscience 30, 477–484.

Millar, J.K., Wilson-Annan, J.C., Anderson, S., Christie, S., Taylor, M.S., Semple, C.A.M., Devon, R.S., Clair, D.M.S., Muir, W.J., Blackwood, D.H.R., et al. (2000). Disruption of two novel genes by a translocation co-segregating with schizophrenia. Human Molecular Genetics 9, 1415–1423.

Miller, R.F. (2004). D-Serine as a glial modulator of nerve cells. Glia 47, 275–283.

Minassian, A., Henry, B.L., Young, J.W., Masten, V., Geyer, M.A., and Perry, W. (2011). Repeated assessment of exploration and novelty seeking in the human behavioral pattern monitor in bipolar disorder patients and healthy individuals. PloS one 6, e24185.

Nagel, G., Szellas, T., Huhn, W., Kateriya, S., Adeishvili, N., Berthold, P., Ollig, D., Hegemann, P., and Bamberg, E. (2003). Channelrhodopsin-2, a directly light-gated cation-selective membrane channel. Proceedings of the National Academy of Sciences of the United States of America 100, 13940–13945.

Nimmerjahn, A., Mukamel, E.A., and Schnitzer, M.J. (2009). Motor behavior activates Bergmann glial networks. Neuron 62, 400–412.

Niwa, M., Cash-Padgett, T., Kubo, K.I., Saito, A., Ishii, K., Sumitomo, A., Taniguchi, Y., Ishizuka, K., Jaaro-Peled, H., Tomoda, T., et al. (2016). DISC1 a key molecular lead in psychiatry and neurodevelopment: No-More Disrupted-in-Schizophrenia 1. Molecular Psychiatry 21, 1488–1489.

Otte, D.M., Barcena de Arellano, M.L., Bilkei-Gorzo, A., Albayram, O., Imbeault, S., Jeung, H., Alferink, J., and Zimmer, A. (2013). Effects of Chronic D-Serine Elevation on Animal Models of Depression and Anxiety-Related Behavior. PLoS One 8, e67131.

Panatier, A., Theodosis, D.T., Mothet, J.P., Touquet, B., Pollegioni, L., Poulain, D.A., and Oliet, S.H. (2006). Glia-derived D-serine controls NMDA receptor activity and synaptic memory. Cell 125, 775–784.

Paoletti, P., Bellone, C., and Zhou, Q. (2013). NMDA receptor subunit diversity: impact on receptor properties, synaptic plasticity and disease. Nature Reviews Neuroscience 14, 383–400.

Papouin, T., Dunphy, J.M., Tolman, M., Dineley, K.T., and Haydon, P.G. (2017a). Septal Cholinergic Neuromodulation Tunes the Astrocyte-Dependent Gating of Hippocampal NMDA Receptors to Wakefulness. Neuron 94, 840–854 e847.

Papouin, T., Henneberger, C., Rusakov, D.A., and Oliet, S.H.R. (2017b). Astroglial versus Neuronal D-Serine: Fact Checking. Trends in Neurosciences 40, 517–520.

Pelluru, D., Konadhode, R.R., Bhat, N.R., and Shiromani, P.J. (2016). Optogenetic stimulation of astrocytes in the posterior hypothalamus increases sleep at night in C57BL/6J mice. The European journal of neuroscience 43, 1298–1306.

Perea, G., Yang, A., Boyden, E.S., and Sur, M. (2014). Optogenetic astrocyte activation modulates response selectivity of visual cortex neurons in vivo. Nature Communications 5, 3262.

Pogorelov, V.M., Rodriguiz, R.M., Insco, M.L., Caron, M.G., and Wetsel, W.C. (2005). Novelty Seeking and Stereotypic Activation of Behavior in Mice with Disruption of the Dat1 Gene. Neuropsychopharmacology 30, 1818–1831.

Porteous, D.J., Millar, J.K., Brandon, N.J., and Sawa, A. (2011). DISC1 at 10: connecting psychiatric genetics and neuroscience. Trends in Molecular Medicine 17, 699–706.

Porteous, D.J., Thomson, P.A., Millar, J.K., Evans, K.L., Hennah, W., Soares, D.C., McCarthy, S., McCombie, W.R., Clapcote, S.J., Korth, C., et al. (2014). DISC1 as a genetic risk factor for schizophrenia and related major mental illness: response to Sullivan. Molecular Psychiatry 19, 141–143.

Raff, M., Abney, E., Cohen, J., Lindsay, R., and Noble, M. (1983). Two types of astrocytes in cultures of developing rat white matter: differences in morphology, surface gangliosides, and growth characteristics. 3, 1289–1300.

Ramirez, F., Moscarello, J.M., LeDoux, J.E., and Sears, R.M. (2015). Active avoidance requires a serial basal amygdala to nucleus accumbens shell circuit. Journal of Neuroscience 35, 3470–3477.

Reddy, L.F., Lee, J., Davis, M.C., Altshuler, L., Glahn, D.C., Miklowitz, D.J., and Green, M.F. (2014). Impulsivity and Risk Taking in Bipolar Disorder and Schizophrenia. Neuropsychopharmacology 39, 456–463.

Rigoli, L., Lombardo, F., and Di Bella, C. (2011). Wolfram syndrome and WFS1 gene. Clinical genetics 79, 103–117.

Rusnakova, V., Honsa, P., Dzamba, D., Ståhlberg, A., Kubista, M., and Anderova, M. (2013). Heterogeneity of Astrocytes: From Development to Injury – Single Cell Gene Expression. PLOS ONE 8, e69734.

Russo, S.J., and Nestler, E.J. (2013). The brain reward circuitry in mood disorders. Nature Reviews Neuroscience 14, 609–625.

Song, H., Stevens, C.F., and Gage, F.H. (2002). Astroglia induce neurogenesis from adult neural stem cells. Nature 417, 39–44.

South, M., Chamberlain, P.D., Wigham, S., Newton, T., Le Couteur, A., McConachie, H., Gray, L., Freeston, M., Parr, J., and Kirwan, C.B. (2014). Enhanced decision making and risk avoidance in high-functioning autism spectrum disorder. Neuropsychology 28, 222.

Stujenske, J.M., and Likhtik, E. (2017). Fear from the bottom up. Nature Neuroscience 20, 765–767.

Swift, M., and Gorman Swift, R. (2000). Psychiatric disorders and mutations at the Wolfram syndrome locus. Biological Psychiatry 47, 787–793.

Swift, R.G., Sadler, D.B., and Swift, M. (1990). Psychiatric findings in Wolfram syndrome homozygotes. Lancet (London, England) 336, 667–669.

Takata, N., Sugiura, Y., Yoshida, K., Koizumi, M., Hiroshi, N., Honda, K., Yano, R., Komaki, Y., Matsui, K., Suematsu, M., et al. (2018). Optogenetic astrocyte activation evokes BOLD fMRI response with oxygen consumption without neuronal activity modulation. Glia 66, 2013–2023.

Takei, D., Ishihara, H., Yamaguchi, S., Yamada, T., Tamura, A., Katagiri, H., Maruyama, Y., and Oka, Y. (2006). WFS1 protein modulates the free Ca2+ concentration in the endoplasmic reticulum. FEBS letters 580, 5635–5640.

Tan, Z., Liu, Y., Xi, W., Lou, H.-f., Zhu, L., Guo, Z., Mei, L., and Duan, S. (2017). Glia-derived ATP inversely regulates excitability of pyramidal and CCK-positive neurons. Nature Communications 8, 13772.

Toker, L., Mancarci, B.O., Tripathy, S., and Pavlidis, P. (2018). Transcriptomic evidence for alterations in astrocytes and parvalbumin interneurons in subjects with bipolar disorder and schizophrenia. Biological psychiatry 84, 787–796.

Tovote, P., Fadok, J.P., and Lüthi, A. (2015). Neuronal circuits for fear and anxiety. Nature Reviews Neuroscience 16, 317–331.

Tropea, D., Hardingham, N., Millar, K., and Fox, K. (2018). Mechanisms underlying the role of DISC1 in synaptic plasticity. The Journal of physiology 596, 2747–2771.

Vigo, D., Thornicroft, G., and Atun, R. (2016). Estimating the true global burden of mental illness. The Lancet Psychiatry 3, 171–178.

Walf, A.A., and Frye, C.A. (2007). The use of the elevated plus maze as an assay of anxiety-related behavior in rodents. Nature Protocols 2, 322–328.

Wang, A.L., Chao, O.Y., Yang, Y.M., Trossbach, S.V., Müller, C.P., Korth, C., Huston, J.P., and de Souza Silva, M.A. (2019). Anxiogenic-like behavior and deficient attention/working memory in rats expressing the human DISC1 gene. Pharmacology, biochemistry, and behavior 179, 73–79.

Wolosker, H. (2011). Serine racemase and the serine shuttle between neurons and astrocytes. Biochimica et Biophysica Acta (BBA) – Proteins and Proteomics 1814, 1558–1566.

Wolosker, H., Balu, D.T., and Coyle, J.T. (2016). The Rise and Fall of the d-Serine-Mediated Gliotransmission Hypothesis. Trends in Neurosciences 39, 712–721.

Wolosker, H., Panizzutti, R., and De Miranda, J. (2002). Neurobiology through the looking-glass: D-serine as a new glial-derived transmitter. Neurochemistry international 41, 327–332.

Yacubian, J., Gläscher, J., Schroeder, K., Sommer, T., Braus, D.F., and Büchel, C. (2006). Dissociable systems for gain- and loss-related value predictions and errors of prediction in the human brain. The Journal of neuroscience : the official journal of the Society for Neuroscience 26, 9530–9537.

Yamamoto, D.J., Woo, C.-W., Wager, T.D., Regner, M.F., and Tanabe, J. (2015). Influence of dorsolateral prefrontal cortex and ventral striatum on risk avoidance in addiction: A mediation analysis. Drug and Alcohol Dependence 149, 10–17.

Yang, J., Yu, H., Zhou, D., Zhu, K., Lou, H., Duan, S., and Wang, H. (2015). Na+–Ca2+ exchanger mediates ChR2-induced [Ca2+]i elevation in astrocytes. Cell Calcium 58, 307–316.

Ye, F., Kang, E., Yu, C., Qian, X., Jacob, F., Yu, C., Mao, M., Poon, R.Y.C., Kim, J., Song, H., et al. (2017). DISC1 Regulates Neurogenesis via Modulating Kinetochore Attachment of Ndel1/Nde1 during Mitosis. Neuron 96, 1041–1054. e1045.

Yong, V., Yong, F., Olivier, A., Robitaille, Y., and Antel, J.J.J.o.n.r. (1990). Morphologic heterogeneity of human adult astrocytes in culture: Correlation with HLA-DR expression. 27, 678–688.

Zatyka, M., Ricketts, C., da Silva Xavier, G., Minton, J., Fenton, S., Hofmann-Thiel, S., Rutter, G.A., and Barrett, T.G. (2008). Sodium-potassium ATPase 1 subunit is a molecular partner of Wolframin, an endoplasmic reticulum protein involved in ER stress. Hum Mol Genet 17, 190–200.

Zhou, X., Wu, B., Liu, W., Xiao, Q., He, W., Zhou, Y., Wei, P., Zhang, X., Liu, Y., Wang, J., et al. (2021). Reduced Firing of Nucleus Accumbens Parvalbumin Interneurons Impairs Risk Avoidance in DISC1 Transgenic Mice. Neuroscience bulletin 37, 1325–1338.

Zhou, X., Xiao, Q., and Tu, J. (2022). Diverse risk-avoidance behaviors in DISC1 mice are associated with different neuronal firing patterns in BLA neurons. Biochemical and biophysical research communications 587, 107–112.

Zhou, Z., Liu, X., Chen, S., Zhang, Z., Liu, Y., Montardy, Q., Tang, Y., Wei, P., Liu, N., Li, L., et al. (2019). A VTA GABAergic Neural Circuit Mediates Visually Evoked Innate Defensive Responses. Neuron 103, 473–488. e476.

## Reference

Franklin, K., and Paxinos, G. (1997). The Mouse Brain in Stereotaxic Coordinates.

Yang, X., Liu, Q., Zhong, J., Song, R., Zhang, L., and Wang, L. (2020). A simple threat-detection strategy in mice. BMC Biology 18, 93.

