## supplemental figures for "Depolarization of astrocytes in the basolateral amygdala restores WFS1 neuronal activity and rescues impaired risk-avoidance behavior in *DISC1^TM^* mice"

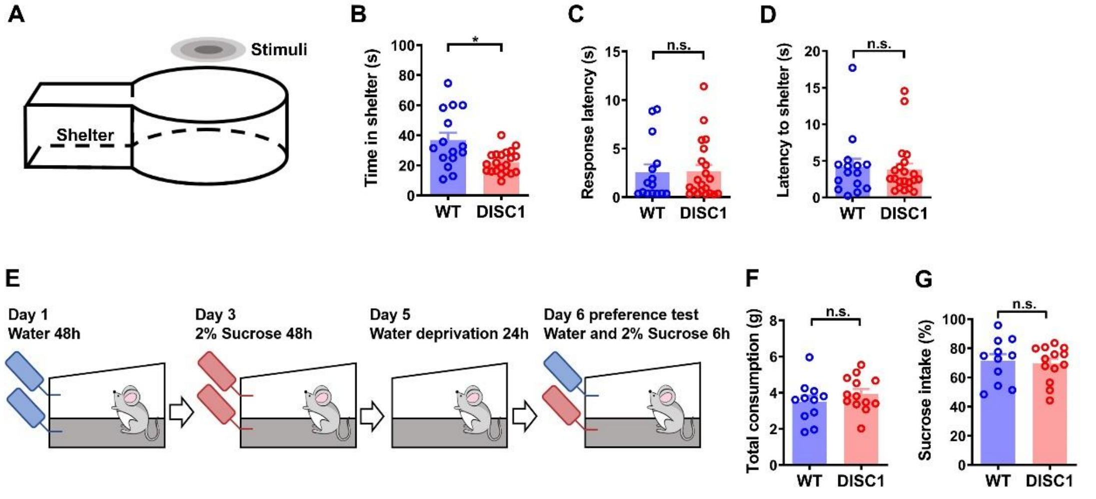


**Supplemental figure 1 The looming test and sucrose preference test of *DISC1-N$^{TM}$* and WT mice**

(A) Schematic showing the looming test.

(B-D) Time spent in shelter (B), response latency following looming stimulation (C) and time taken to return to the shelter (D) during the looming task (unpaired t-test, *$P$ = 0.0124; n =15 from 5 WT mice, n =21 from 7 *DISC1-N$^{TM}$* mice).

(E) Schematic showing the sucrose preference test.

(F-G) Total fluid consumption (F) and sucrose preference (G) during the sucrose preference test (unpaired t-test, n $_{(WT)}$ =11, n $_{(DISC1)}$ =13).

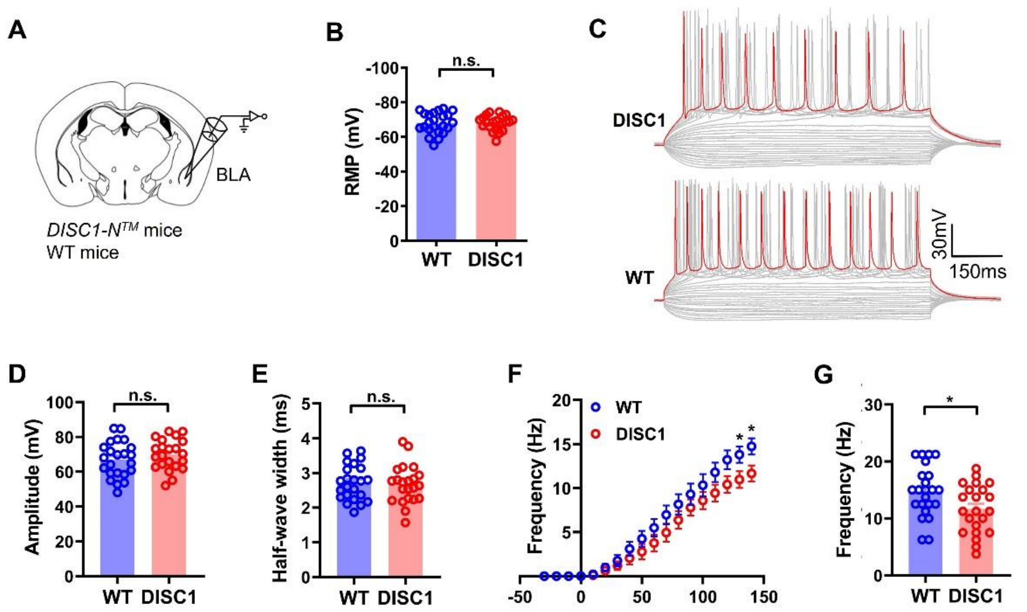


**Supplemental figure 2 Electrophysiological properties of neurons show that the BLA plays a role in risk-avoidance behavior in *DISC1-N$^{TM}$* mice**

(A) Schematic of brain slice patch clamp technique.

(B) Comparison of the RMP of BLA neurons in WT mice and *DISC1-N$^{TM}$* mice (unpaired *t*-test; n =23 neurons from 11 WT mice, n = 22 neurons from 10 *DISC1-N$^{TM}$* mice).

(C) Representative traces of BLA neurons in response to current injection (from -100 pA to +140 pA, 10 pA interval, red traces represent 140 pA current injection)

(D-E) Comparison of the amplitude (E) and half-wave width (F) of action potentials generated by BLA neurons when stimulated by 140 pA current (unpaired t- test; n =23 neurons from 11 WT mice, n = 22 neurons from 10 *DISC1-N$^{TM}$* mice).

(F) Current stimulation-firing frequency curve of BLA neurons from WT and *DISC1-N$^{TM}$* mice (unpaired t-test, *$P$ (left) = 0.0375, *$P$ (right) = 0.0228; n = 23 neurons from 11 WT mice, n = 22 neurons from 10 *DISC1-N$^{TM}$* mice).

(G) The frequency of action potentials generated by neurons in the BLA when stimulated by 140 pA current (unpaired *t*-test, *$P$ = 0.0228; n = 23 neurons from 11 WT mice, n = 22 neurons from 10 *DISC1-N$^{TM}$* mice).

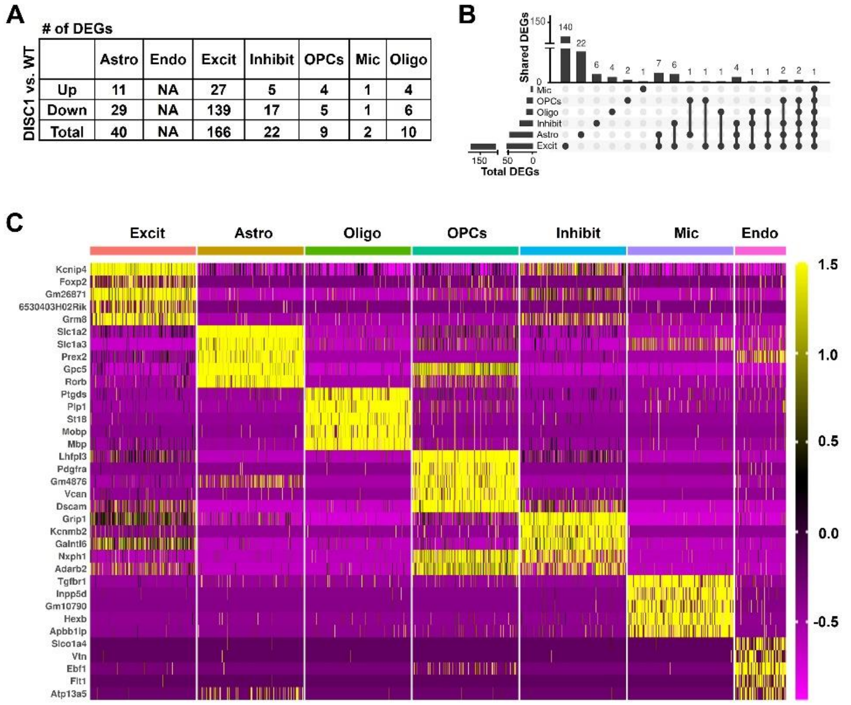


**Supplemental figure 3 Statistics of DEGs and enriched genes from snRNA-seq analysis of the BLA**

(A) The numbers of DEGs between DISC1 and WT samples within each cell type (adjusted $P < 0.1$, log2 fold change $\geq 0.1$ or $\leq -0.1$). Down: down-regulated; Up: up-regulated.

(B) Upset plot showing the coregulated DEGs among all six cell types.

(C) Heatmap showing the top five most enriched genes for each cell type.

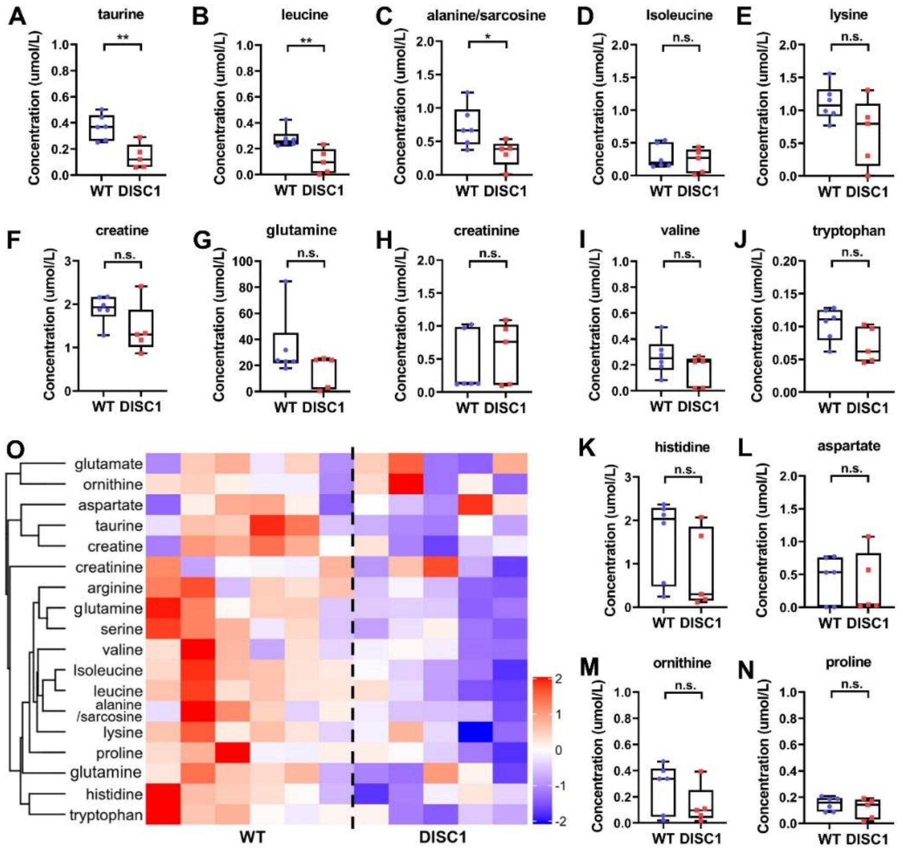


**Supplemental figure 4 LC-MS analysis of BLA dialysate from *DISC1-N*$^{TM}$ and WT mice**

(A-N) The partial results of LC-MS analysis of the BLA dialysate from WT mice and *DISC1-N*$^{TM}$ mice (unpaired t-test, from left to right: $^{**}P$ = 0.0039 and 0.007, $^{*}P$ = 0.0364; $n_{(WT)}$ =6, $n_{(DISC1)}$ =5).

(O) Heatmap display of the hierarchical clustering results of some amino acids detected by LC-MS.

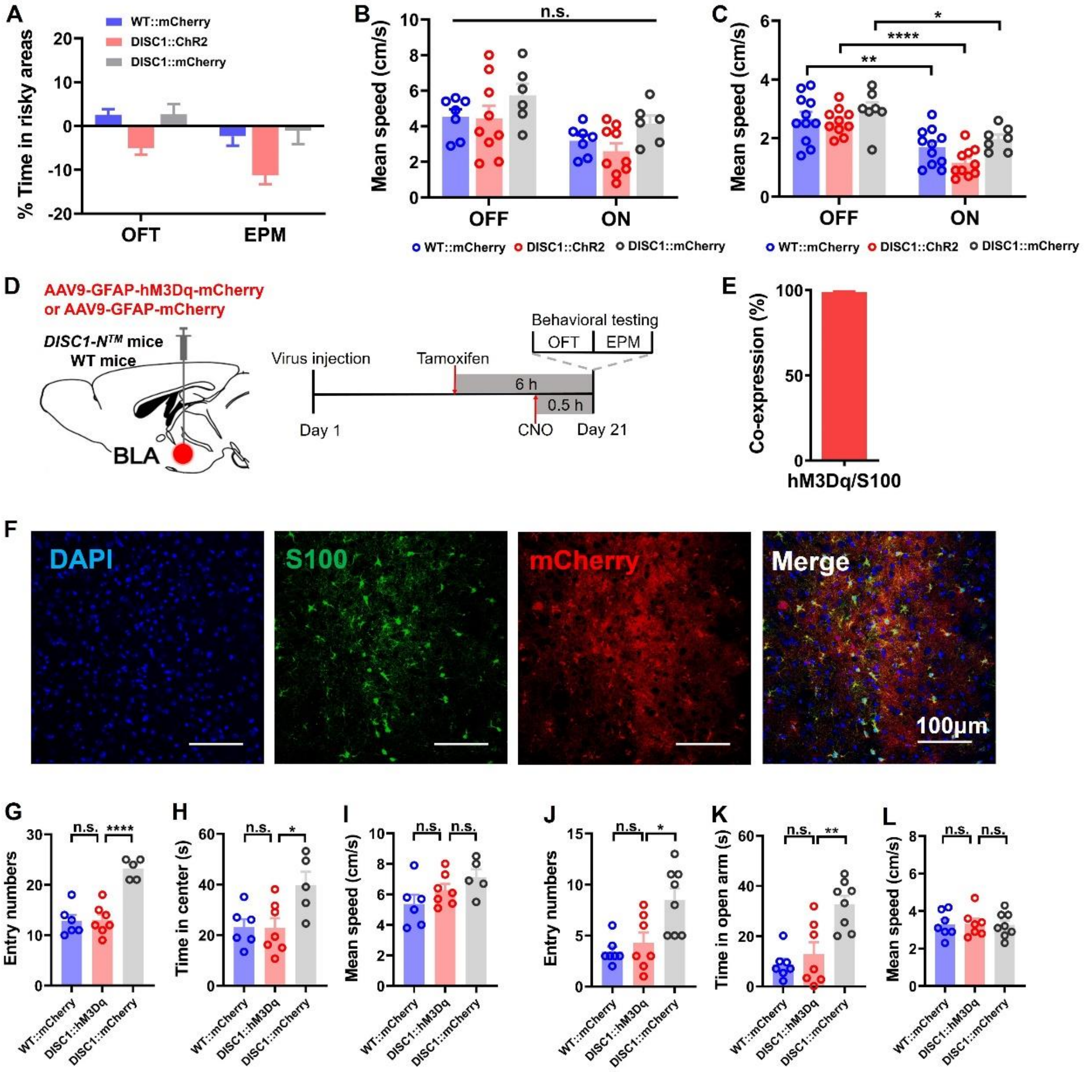


**Supplemental figure 5 Pharmacogenetic tools to modulate BLA astrocytes in *DISC1-N^TM* mice**

(A) The percentage display of time spent in the risky areas during the OFT during the optogenetic stimulation paradigms (Figure 3D-I).

(B-C) The mean speed during the OFT (B) and EPM test (C) during the optogenetic stimulation paradigms.

(D) Schematic showing the experimental process.

(E) The percentage of co-expression of hM3Dq-expressing cells and S100-expressing cells in the BLA ($n_{(DISC1)} = 3$).

(F) Astrocytes in the BLA infected with AAV-GFAP-hM3Dq-mCherry (red) co-stained with S100 (green).

(G-I) The number of entries to (G), and time spent in the center (H) and the mean speed (I) after

CNO administration during OFT (unpaired t-test, $^{*}P = 0.0233$, $^{****}P < 0.0001$; $n_{(WT\text{-}mCherry)} = 6$, $n_{(DISC1\text{-}hM3Dq)} = 7$, $n_{(DISC1\text{-}mCherry)} = 5$).

(J-L) Number of entries to (J), and time spent in the open arms (K) and the mean speed (L) after CNO administration during the EPM test (unpaired t-test, $^{*}P = 0.0171$, $^{**}P = 0.0041$; $n_{(WT\text{-}mCherry)} = 7$, $n_{(DISC1\text{-}hM3Dq)} = 7$, $n_{(DISC1\text{-}mCherry)} = 8$).

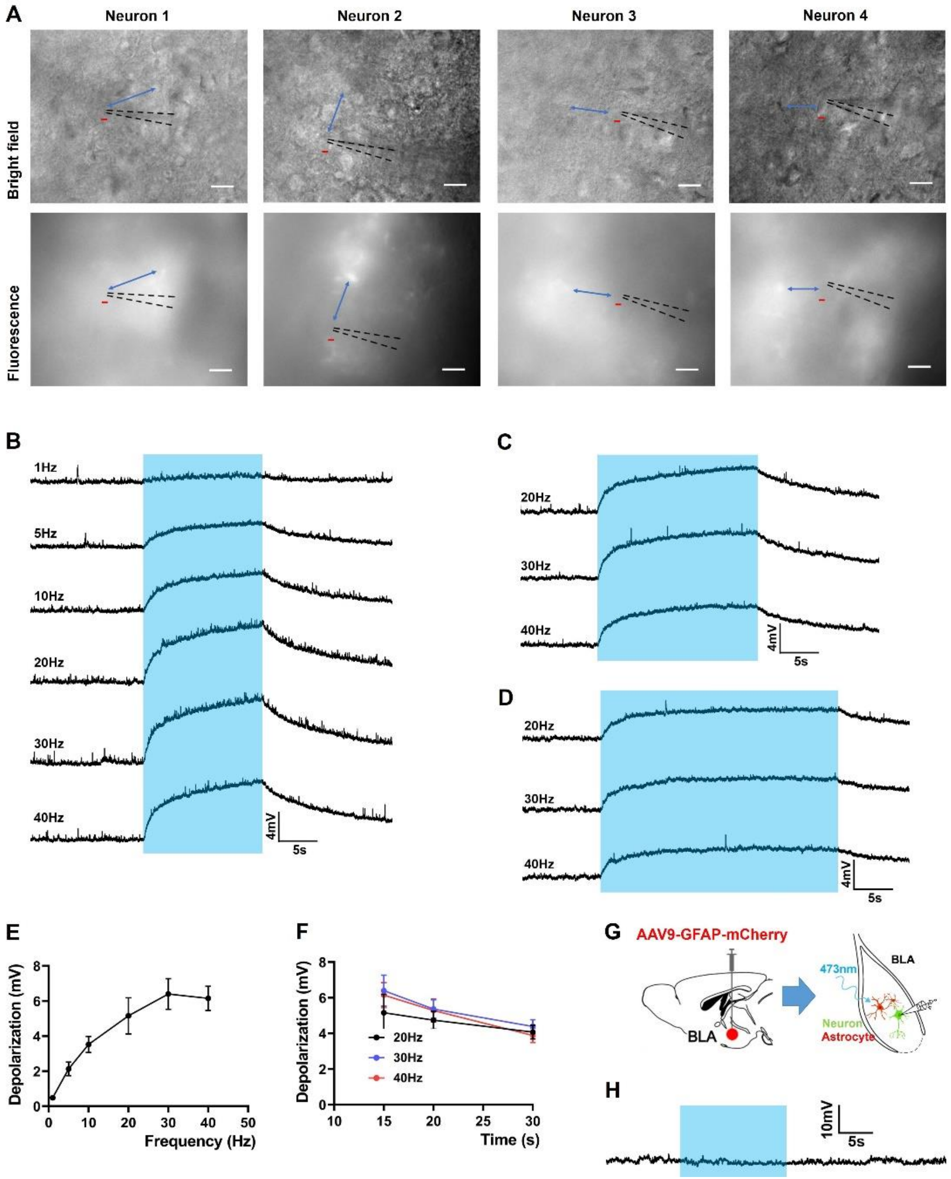


**Supplemental figure 6 Electrophysiological responses of low-response neurons to different patterns of light stimulation**

(A) Representative images showing the distance (blue bidirectional arrows) between the patch clamp recording sites (red short line) and the brightest fluorescent (mCherry) expression sites (black dotted lines represent recording electrodes, scale bars indicate 40 µm).

(B-D) Changes in membrane potential of low-response neurons under light stimulation of different frequencies for 15 s (B), 20 s (C) and 30 s (D).

(E) The depolarization of membrane potential in low-response neurons during 15 s light stimulation under different frequencies (n = 6 neurons from 4 WT mice).

(F) The depolarization of membrane potential in low-response neurons at 20 Hz, 30 Hz and 40 Hz frequency under different durations (n = 6 neurons from 4 WT mice).

(G) Schematic showing patch clamp recording of neurons in the BLA.

(H) Representative trace showing no change in membrane potential of nearby neurons when blue light was shone on astrocytes that do not express ChR2.

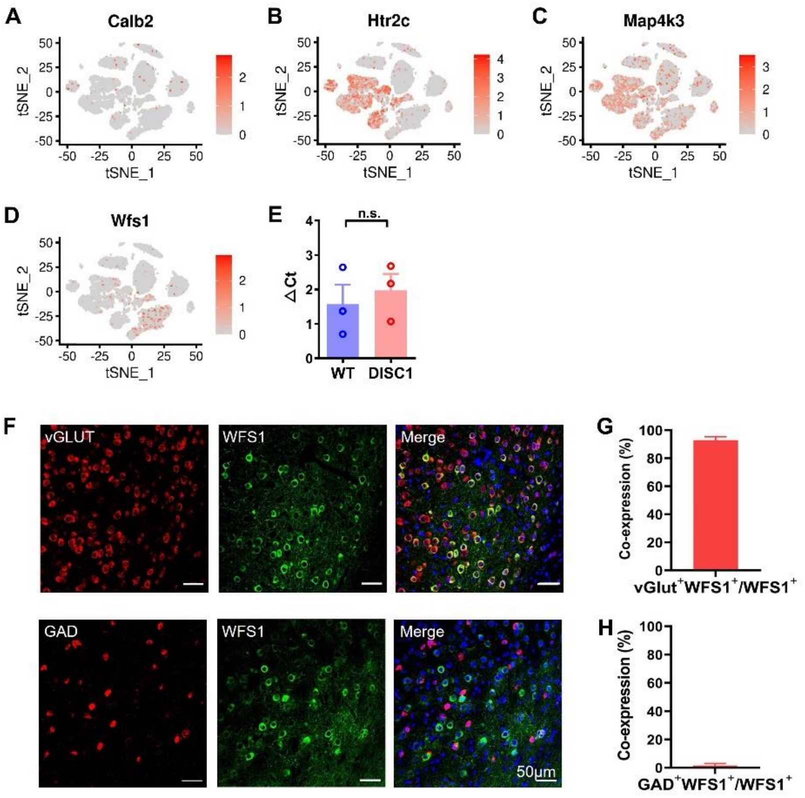


**Supplemental figure 7 WFS1 neurons in the BLA are excitatory neurons**

(A-D) The t-SNE plots showing the distributions of the four neuronal markers in the $DISC1\text{-}N^{TM}$ BLA samples for snRNA-seq.

(E) The expression of *WFS1* mRNA in $DISC1\text{-}N^{TM}$ and WT mice (unpaired t-test, $n_{(WT)}=3$, $n_{(DISC1)}=3$).

(F) Representative images showing co-expression of WFS1 neurons in the BLA stained with vGlut (top) and GAD (bottom).

(G) Quantification of the proportion of neurons expressing vGlut and WFS1 ($n_{(WT)}=3$).

(H) Quantification of the proportion of neurons expressing GAD and WFS1 ($n_{(WT)}=3$).
